# A conserved and tunable mechanism for the temperature-controlled condensation of the translation factor Ded1p

**DOI:** 10.1101/2022.10.11.511767

**Authors:** Ceciel Jegers, Titus M. Franzmann, Julian Hübner, Jakob Schneider, Cedric Landerer, Sina Wittmann, Agnes Toth-Petroczy, Remco Sprangers, Anthony A. Hyman, Simon Alberti

## Abstract

Heat shock promotes the assembly of translation factors into condensates to facilitate the production of stress-protective proteins. How translation factors detect heat and assemble into condensates is not well understood. Here, we investigate heat-induced condensate assembly by the translation factor Ded1p from five different fungi, including Ded1p from *Saccharomyces cerevisiae*. Using targeted mutagenesis and *in vitro* reconstitution biochemistry, we find that heat-induced Ded1p assembly is driven by a conformational rearrangement of the folded helicase domain. This rearrangement determines the assembly temperature and the assembly of Ded1p into nanometer-sized particles, while the flanking intrinsically disordered regions engage in intermolecular interactions to promote assembly into micron-sized condensates. Using protein engineering, we identify six amino acid substitutions that determine most of the thermostability of a thermophilic Ded1p ortholog, thereby providing a molecular understanding underlying the adaptation of the Ded1p assembly temperature to the specific growth temperature of the species. We conclude that heat-induced assembly of Ded1p into translation factor condensates is regulated by a complex interplay of the structured domain and intrinsically disordered regions which is subject to evolutionary tuning.

## Introduction

Temperature affects all aspects of life by setting the pace for cellular processes and biochemical reactions. Proteins are among the most abundant macromolecules in the cell, and temperature-induced changes to the native structure of proteins pose a serious threat to cell survival. With increasing temperature, proteins are prone to unfolding and aggregation, and key proteins lose activity, ultimately leading to heat-induced cell death (Jarzab et al., 2020; Leuenberger et al., 2017).

To survive heat stress, cells selectively increase the expression of stress-protective genes. This phenomenon is known as the heat shock response (HSR) and serves to reduce the burden of protein misfolding and aggregation (Lindquist, 1986; Lindquist and Craig, 1988). The HSR is vital for sessile and ectothermic organisms, such as *Saccharomyces cerevisiae*, a budding yeast from the Saccharomycetes class and a member of the Saccharomyceta clade of fungi*. S. cerevisiae* survives exposure to temperatures between 3°C and 45°C, and exhibits optimum growth near 32°C (Salvado et al., 2011). However, a five-degree increase from 32°C to 37°C already activates the HSR (Verghese et al., 2012).

The HSR is highly conserved and the temperature at which it is activated is fine-tuned to the thermal niche of the organism. For example, the proteomes of thermophilic organisms are more resilient to heat-induced unfolding than the proteomes of mesophilic organisms (Jarzab et al., 2020; Leuenberger et al., 2017) and, accordingly, their HSR is typically activated at higher temperatures (Fields, 2001; Somero, 1995). What determines the thermostability of proteins has been studied with various model proteins. These studies revealed that temperature adaptation of proteins generally does not require large structural changes but can be mediated by few amino acid substitutions that, for instance, introduce (de)stabilizing interactions and/or alter molecular packing (Somero, 1995; Taylor and Vaisman, 2010).

While most proteins exhibit unfolding transitions above the viable temperature of an organism, some proteins are only marginally stable and unfold within the physiological temperature range of an organism (Wallace et al., 2015). Examples are several translation factors, which assemble into higher-order structures and change from the soluble to the insoluble fraction in budding yeast exposed to sublethal temperatures (Cherkasov et al., 2015; Iserman et al., 2020; Kroschwald et al., 2018; Riback et al., 2017; Wallace et al., 2015). The fact that these translation factors carry out essential functions in protein synthesis in growing cells suggests that their heat-induced assembly into condensates upon heating may not be harmful, but an adaptative mechanism to sense environmental changes and regulate translation (Wallace et al., 2015, Franzmann and Alberti, 2019a; Yoo et al., 2019).

The RNA helicase Ded1p is an example of a protein that reversibly assembles into translation factor condensates in the yeast cytoplasm upon exposure to heat. This essential DEAD-box helicase is one of the first proteins to condense upon exposure to heat stress (Iserman et al., 2020; Wallace et al., 2015) and this process suppresses the translation of mRNAs that encode housekeeping proteins (Iserman et al., 2020; Sen et al., 2021). Concomitantly, the sequestration of Ded1p has been proposed to promote the preferential production of stress-protective proteins, thereby complementing the transcriptional arm of the HSR with translational regulation. *In vitro* reconstitution experiments revealed that Ded1p autonomously assembles into condensates upon heating by the process of phase separation (Iserman et al., 2020). The underlying molecular mechanisms underlying this adaptive response have remained largely undefined.

Here, we investigated the molecular mechanism of heat-induced Ded1p condensation and asked if and how the Ded1p assembly temperature is adapted to the temperature niche of organisms. To this end, we identified Ded1p orthologs from different fungi and quantitatively characterized the temperature behavior. We demonstrate that the ability of Ded1p to form condensates upon heating is conserved among the fungal orthologs and that the onset temperature for assembly is adapted to the growth temperature of the respective species. Adaptation of the Ded1p assembly temperature requires evolutionary tuning of the structured helicase domain, and the assembly temperature is determined by the structural stability of the helicase domain. Mutational and structural analyses revealed that the IDRs interact with the helicase domain and that the N-terminal IDR stabilizes the native helicase domain fold of Ded1p. We further show that the IDRs engage in intermolecular protein interactions upon heating and support assembly into micron-sized condensates. We propose that heat-induced assembly of translation factors involves a complex interplay between structured domains and IDRs, which determines both the thermal responsiveness of the protein and its ability to control the translational HSR. We suggest that translation factor assembly into condensates is adapted to the specific thermal niche of an organism and plays a vital role in the execution of the HSR.

## Results

### Identification of Ded1p orthologs from species living in different thermal niches

Ded1p is a DEAD-box helicase that uses ATP binding and hydrolysis to unwind complex secondary structures in RNA and to promote translation initiation in yeast (Iost et al., 1999). Upon heat shock, Ded1p assembles into translation factor condensates (Iserman et al., 2020; Wallace et al., 2015). Its sequestration into condensates has been linked to a decrease in the translation of housekeeping and the preferential synthesis of stress protective proteins (Iserman et al., 2020; Sen et al., 2021). While being critical for the cellular response to heat stress, the mechanism by which Ded1p detects changes in environmental temperature and assembles into stress-protective condensates remained unknown.

To provide a mechanistic understanding of heat-induced condensation of Ded1p, we set out to identify Ded1p orthologs from different fungi and characterize their behavior upon temperature increases. The availability of a large collection of fungal genomes combined with data on the preferred growth conditions of fungi provided an opportunity to identify Ded1p orthologs and determine how temperature shapes the evolution of Ded1p sequences. We identified a set of 630 orthologous Ded1p sequences from genome analysis and found that most of the identified Ded1p orthologs clustered phylogenetically according to their fungal class (**Supplemental Figure 1**). To identify Ded1p orthologs with different temperature profiles, we selected yeast species from different temperature climates from the Saccharomycetes and Sordariomycetes classes (**Figure 1A** and **Supplemental Figure 1**).

**Figure 1.**
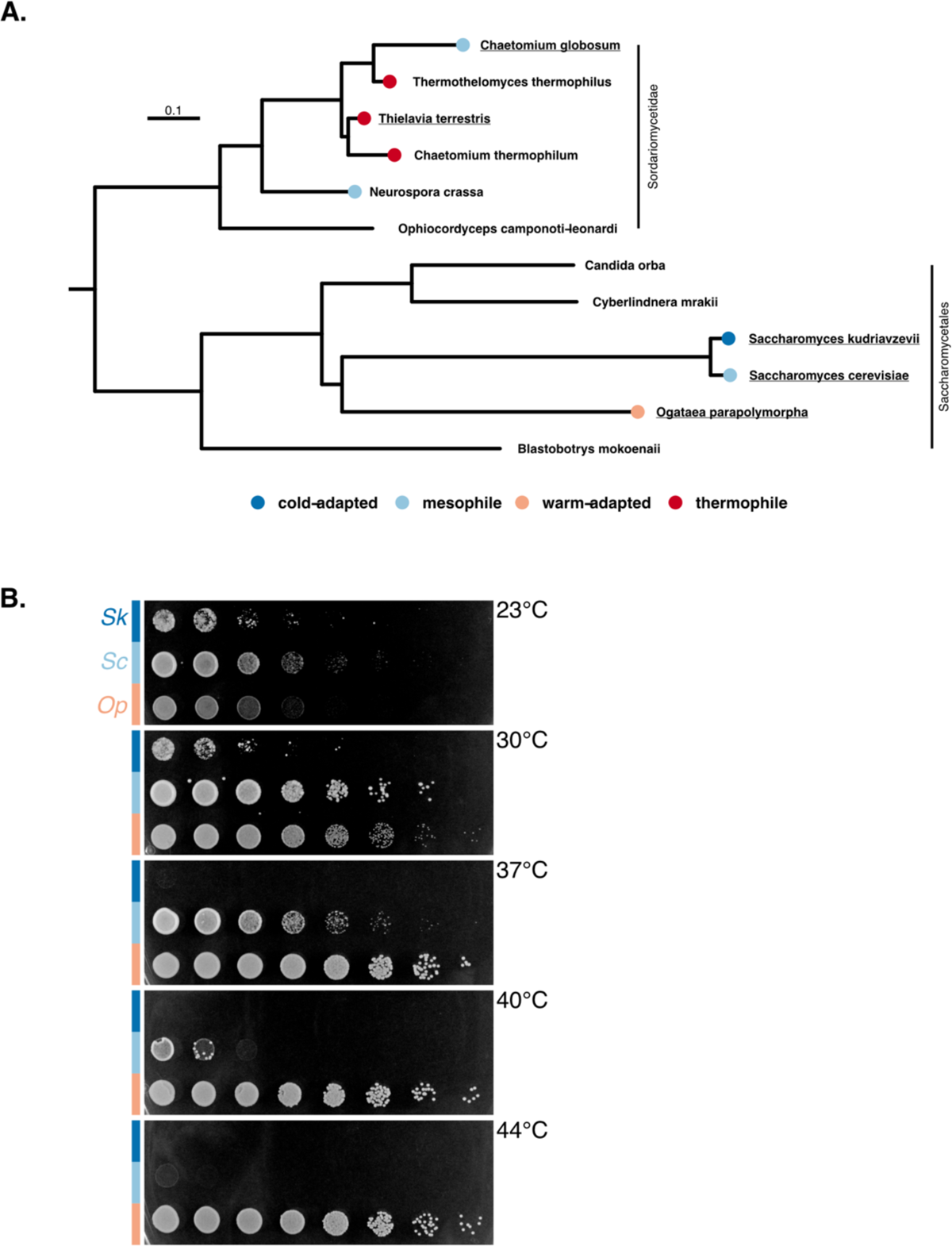
Identification of Ded1p orthologs from species with different growth temperatures. **A.** Phylogenetic tree based on fungal Ded1p orthologs from selected Sordariomycetes and Saccharomycetes species. Colors refer to the growth temperature classification of the respective species (see text). Scale = 0.1 substitutions per site. **B.** Yeast spotting assay with 5-fold serial dilutions of *S. kudriavzevii* (*Sk*), *S. cerevisiae* (*Sc*) and *O. parapolymorpha* (*Op*) grown on YPD at the indicated temperatures for 24 h.

These classes contain budding yeast (Saccharomycetes) and filamentous fungi (Sordariomycetes). From the Saccharomycetes class, we selected *Saccharomyces kudriavzevii (Sk), Saccharomyces cerevisiae (Sc)* and *Ogataea parapolymorpha (Op)*. From the Sordariomycetes class, we selected *Chaetomium globosum (Cg)* and *Thielavia terrestris (Tt)*.

Growth assays at different temperatures revealed that *Sk* did not grow at T ≥ 37°C and *Sc* did not grow at T ≥ 44°C after 24 h (**Figure 1B**). *Op* grew at all tested temperatures (**Figure 1B**). These differences in growth temperature are consistent with published data and suggest that, relative to the mesophile *Sc*, the growth temperature of *Sk* is cold-adapted (Salvado et al., 2011) and the growth of *Op* is warm-adapted. Regarding the species from the Sordariomycetes class, *Cg* has previously been shown to grow well at 34°C but not at ≥45°C, and *Tt* grew even at 55°C, which contributed to their classification as meso- and thermophile respectively (Morgenstern et al., 2012). In summary, we identified the sequences of Ded1p orthologs from fungi from different classes and these fungi exhibit different temperature growth profiles. Based on the difference in growth temperature, we speculate that the Ded1p orthologs from these species function at different temperature ranges and a quantitative comparison of the temperature response of these orthologs can provide a molecular understanding for heat-induced condensate assembly.

### The species’ growth temperature correlates with the heat-induced Ded1p assembly

Previous data using budding yeast suggested that heat induces Ded1p condensation to regulate the translation of housekeeping mRNAs (Iserman et al., 2020). To test if Ded1p orthologs exhibit different assembly temperatures compared to Ded1p from *S. cerevisiae*, we expressed and purified *Sc* Ded1p and orthologs as C-terminal GFP-tagged fusion proteins from insect cells and characterized them *in vitro*. Consistent with published data (Iserman et al., 2020), *Sc* Ded1p assembled into reversible, spherical and amorphous condensates in the presence or absence of RNA, respectively (**Figure 2A**). Despite morphological differences, assemblies of *Sc* Ded1p with and without RNA were detected by microscopy at ∼38°C (**Figure 2A**) (Iserman et al., 2020) (Note: Subsequent experiments were conducted in the absence of RNA).

**Figure 2.**
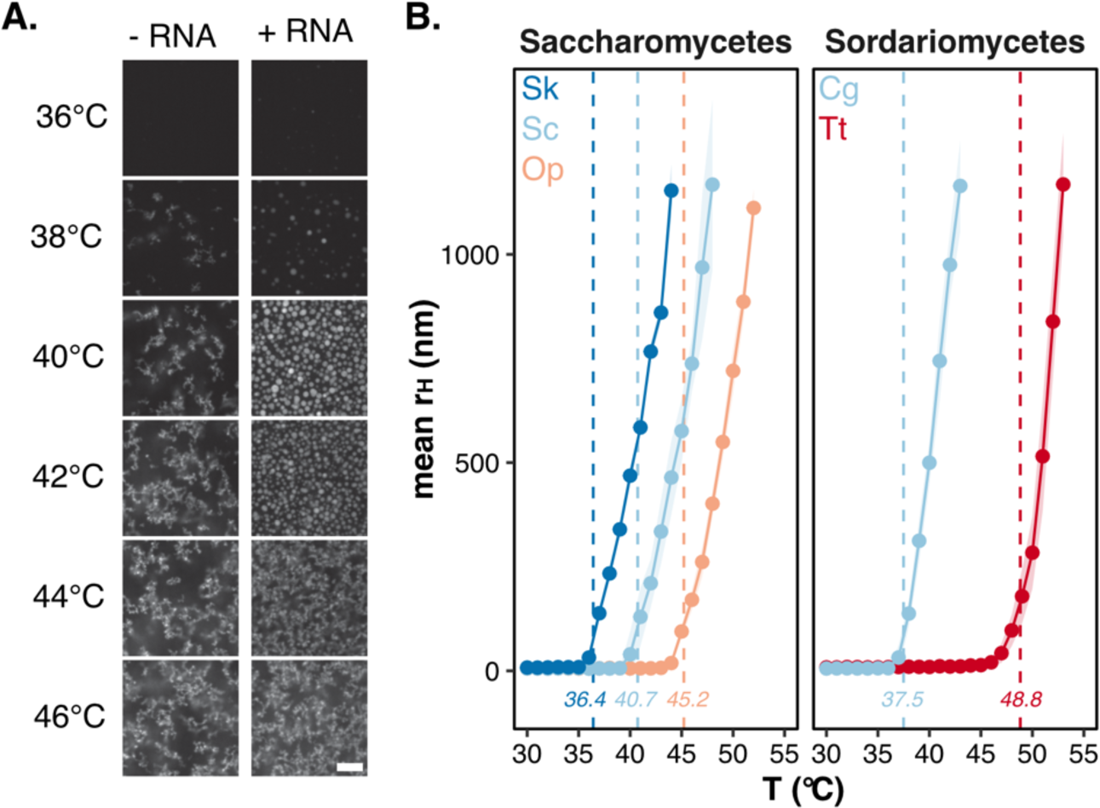
The assembly temperature of Ded1p is species-specific. **A.** Representative fluorescence microscopy images of GFP-labelled *S. cerevisiae* Ded1p heated to indicated temperatures in absence (-RNA) and presence (+RNA) of *in vitro* transcribed RNA. Scale bar = 10 μm. **B.** Hydrodynamic radius (r_H_) as a function of temperature for GFP-labelled Ded1p orthologs from budding yeast (*S. cerevisiae* (Sc), *S. kudriavzevii* (Sk)*, O. parapolymorpha* (Op), left) and filamentous fungi (*C. globosum* (Cg) and *T. terrestris* (Tt), right) using DLS. Assembly T_onset_ for each protein is highlighted with a dashed line. Mean (points), sd (light ribbon), n = 3-4.

Next, we determined the assembly temperatures for *Sc* Ded1p and the orthologs from *Sk, Op, Cg* and *Tt,* complementing the dataset from Iserman et al. (2020) with additional Ded1p orthologs. First, we monitored the hydrodynamic radius (r_H_) as a function of temperature using dynamic light scattering. At 20°C, the r_H_ of the most abundant protein species was similar among Ded1p orthologs (*Sc* = 6.0 nm ± 0.7 nm, *Sk* = 6.2 nm ± 0.3 nm, *Op* = 5.9 nm ± 0.5 nm, *Cg* = 5.7 nm ± 0.1 nm and *Tt* = 5.9 nm ± 0.4 nm). Upon temperature increase, the mean r_H_ remained relatively constant until a protein-specific temperature (**Figure 2B**) at which the r_H_ increased, indicating assembly into larger particles. Inspecting these samples by fluorescence microscopy revealed that all proteins assembled into morphologically similar condensates (**Supplemental Figure 2A-B**). Compared to Ded1p from mesophilic *Sc,* the Ded1p ortholog from the cold-adapted *Sk* assembled at a lower temperature (ΔT = −4.3°C) and the ortholog from the warm-adapted *Op* at a higher temperature (ΔT = 4.5°C) (**Figure 2B, Table 1)**. Similarly, the T_onset_ of the Ded1p ortholog from the mesophilic Sordariomycetes species *Cg* was 11.3°C below that of the closely related thermophilic ortholog from *Tt* (**Figure 2B**). Comparison of the assembly temperatures and yeast upper growth temperatures (**Table 1**) revealed that that the assembly temperature of Ded1p orthologs correlated and scaled with the respective species’ upper growth temperature.

**Table 1.**
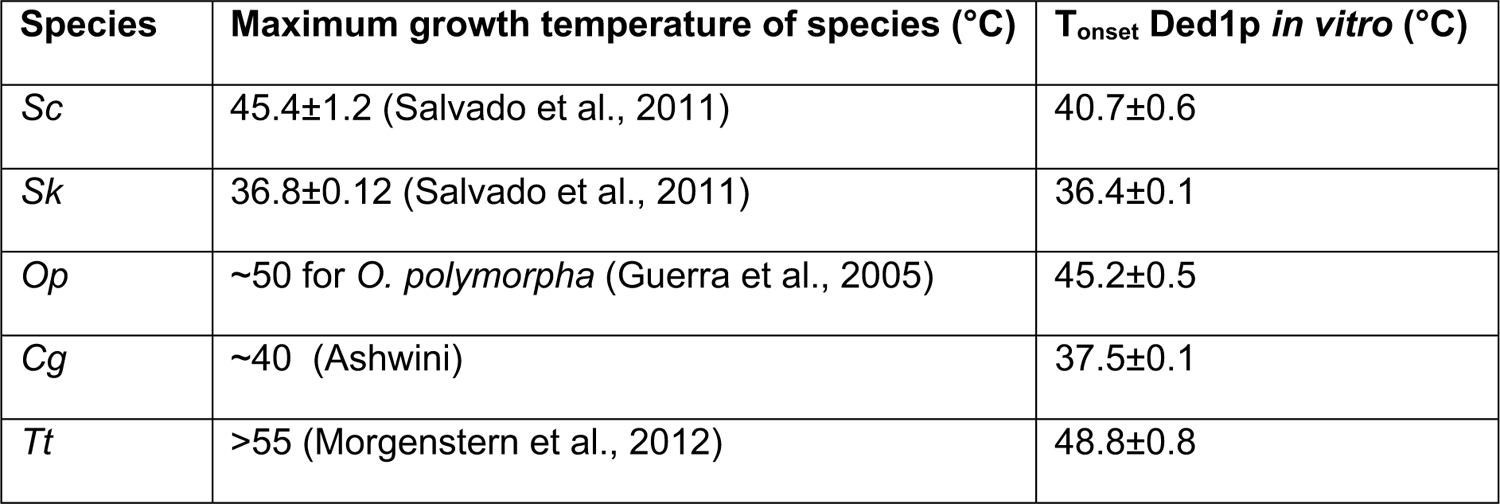
The maximum growth temperatures of selected fungi are listed, as well as the onset temperature for assembly of the respective Ded1p ortholog in vitro.

| Species | Maximum growth temperature of species (°C) | $T_{onset}$ Ded1p *in vitro* (°C) |
| --- | --- | --- |
| *Sc* | 45.4±1.2 (Salvado et al., 2011) | 40.7±0.6 |
| *Sk* | 36.8±0.12 (Salvado et al., 2011) | 36.4±0.1 |
| *Op* | ~50 for *O. polymorpha* (Guerra et al., 2005) | 45.2±0.5 |
| *Cg* | ~40 (Ashwini) | 37.5±0.1 |
| *Tt* | >55 (Morgenstern et al., 2012) | 48.8±0.8 |

### The heat-induced assembly of Ded1p orthologs coincides with a structural transition of the helicase domain

Heat-induced assembly of *Sc* Ded1p coincides with tertiary structure changes (Iserman et al., 2020). To test if the assembly of Ded1p orthologs is also accompanied by changes in tertiary structure, we recorded fluorescence intensities at 350 and 330 nm as a function of temperature. To distinguish structure changes from temperature-induced fluorescence quenching, we analyzed the fluorescence emission ratio (F350/F330) (**Supplemental Figure 3A**). The fluorescence temperature profiles were characterized by a sigmoid shape with an initial plateau with a drift (native baseline) preceding a steep increase (transition) followed by a final plateau with a drift (unfolded baseline) (**Supplemental Figure 3D**). The sigmoid change in fluorescence emission ratio is characteristic of cooperative (un)folding transition and demonstrates that all Ded1p orthologs adopted a stable fold at temperatures below the ortholog specific transition temperature. To determine the transition onset (T_onset_) and the transition midpoints (T_m_), we fitted the data to a two-state transition model. For clarity, the transition profiles are represented as normalized transition data (**see methods for details**). The T_onset_ for the structural change was smallest for Ded1p ortholog from the cold-adapted *Sk* (39.5°C ±0.2°C), followed by the ortholog from the mesophile *Sc* (44.3°C ±0.1°C) and the ortholog from the warm-adapted *Op* (46.3°C ±0.2°C) (**Figure 3A**). Similarly, the orthologs from the mesophile *Cg* and the thermophile *Tt* underwent structural changes at 38.2°C ±0.2°C and at 51.4°C ±0.2°C, respectively (**Figure 3A**). Importantly, the unfolding transition coincided with an increase in light scattering (**Figure 3A, Supplemental Figure 3E**), suggesting that the heat-induced structural change coincided with condensate assembly for all Ded1p orthologs.

**Figure 3.**
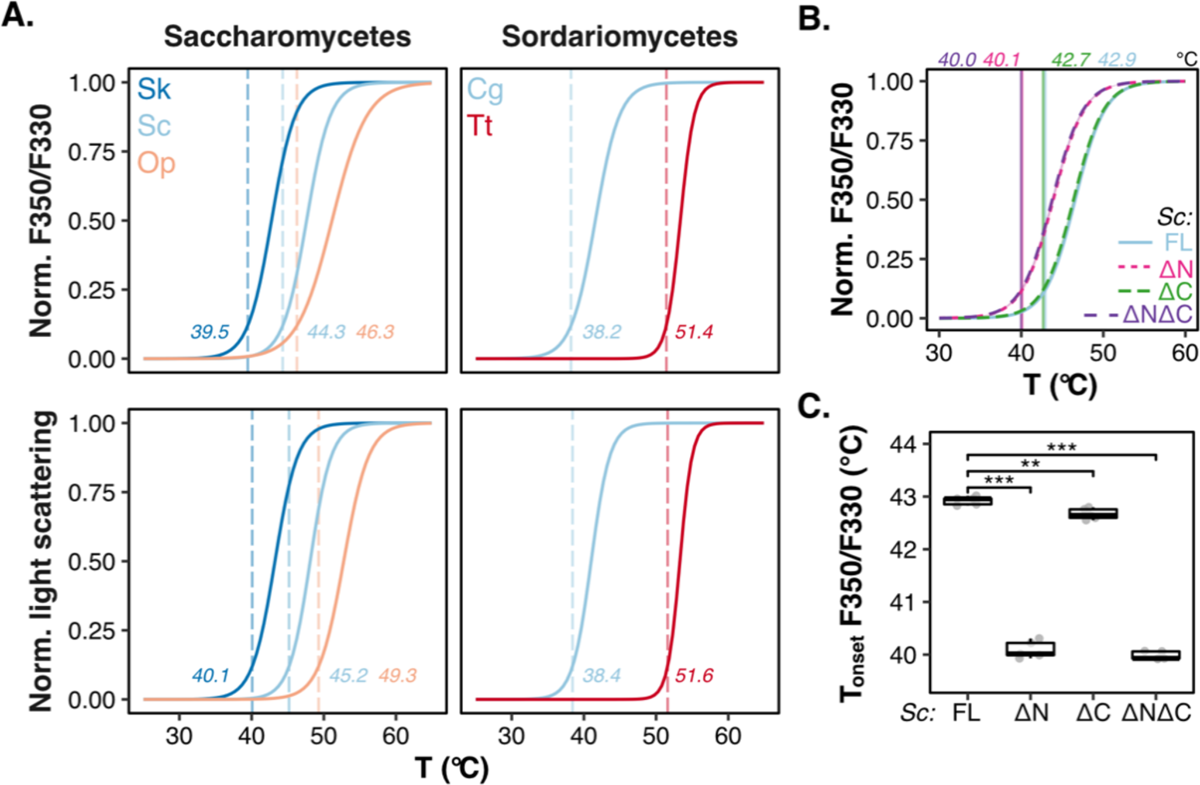
Heat-induced assembly of Ded1p orthologs coincides with a structural transition of the helicase domain. **A.** Change in normalized F350/F330 (top) and back reflection light scattering (bottom) as a function of temperature for 5 μM GFP-labelled Ded1p orthologs from *S. cerevisiae* (Sc), *S. kudriavzevii* (Sk)*, O. parapolymorpha* (Op), *C. globosum* (Cg) and *T. terrestris* (Tt) (bottom). A trendline is shown as a guide and the T_onset_ is highlighted with a dashed line. Mean (solid lines), sd (light ribbon), n = 6. **B.** Change in normalized F350/F330 as a function of temperature for 6 μM GFP-labelled full-length *S. cerevisiae* Ded1p (Sc-FL), Ded1p-βN, -βC and -βNβC using nanoDSF. A trendline is shown as a guide and the T_onset_ is highlighted. Mean (solid line), sd (light ribbon), n = 5. **C.** Apparent T_onset_ of F350/F330 for 6 μM full-length GFP-labelled *S. cerevisiae* Ded1p (Sc-FL), Ded1p-βN, -βC and -βNβC using nanoDSF (n = 5, *** p < 0.001 calculated using T-test).

The heat-induced cooperative increase in fluorescence ratio (**Figure 3A**) suggested that the local environment of one or more tryptophan residues within the Ded1p orthologs changes during unfolding (**Figure 3A**). Full-length *Sc* Ded1p contains seven tryptophan residues, which are distributed throughout its primary structure (**Supplemental Figure 3B**). The tryptophan residue W253 is conserved and located in the globular RecA1 domain, which together with the RecA2 domain constitutes the helicase domain (**Supplemental Figure 3B-C**). Other tryptophan residues in Ded1p are less well conserved and reside in the disordered N- and C-terminal regions that flank the helicase fold (**Supplemental Figure 3B**).

To determine the contributions of tryptophan residues located in different Ded1p domains to the heat-induced fluorescence change, we generated *Sc* Ded1p deletion variants lacking either the N-terminal (ΔN), C-terminal (ΔC) or both IDRs (ΔNΔC) and recorded fluorescence temperature profiles (**Figure 3B**). Like full-length Ded1p (FL), all IDR deletion variants exhibited a cooperative (un)folding transition (**Figure 3B**), indicating that the changes in fluorescence originate from a structural transition of the helicase domain. It is interesting to note that the onset temperature of the ΔC variant was comparable to that of the wildtype protein (**Figure 3C**), indicating that the C-terminal IDR does not affect the stability of the helicase domain much. However, the deletion variants ΔN and ΔNΔC exhibited lower onset temperatures compared to that of the full-length *Sc* Ded1p (**Figure 3C**), suggesting that the N-terminal IDR stabilizes the overall stability of the helicase domain (see below).

Taken together our data suggest that a cooperative structural transition of the helicase domain coincides with Ded1p assembly, and that the assembly temperature is adapted to the physiological growth temperature of the respective yeast species (**Figure 3A**). To provide further evidence that the Ded1p assembly temperature is determined by the structural stability of the helicase domain, we determined the onset temperatures of *Sc* Ded1p in the presence of the destabilizing denaturant guanidine hydrochloride (GdnHCl) or the stabilizing co-solvent glycerol. Within the tested concentration ranges, the apparent unfolding and assembly temperatures of Ded1p decreased linearly as a function of increasing GdnHCl concentrations (**Figure 4**) and increased linearly with increasing glycerol concentrations (**Figure 4**) and remained correlated across the tested ranges. This suggested that the unfolding of the helicase domain determines the assembly of Ded1p into condensates.

**Figure 4.**
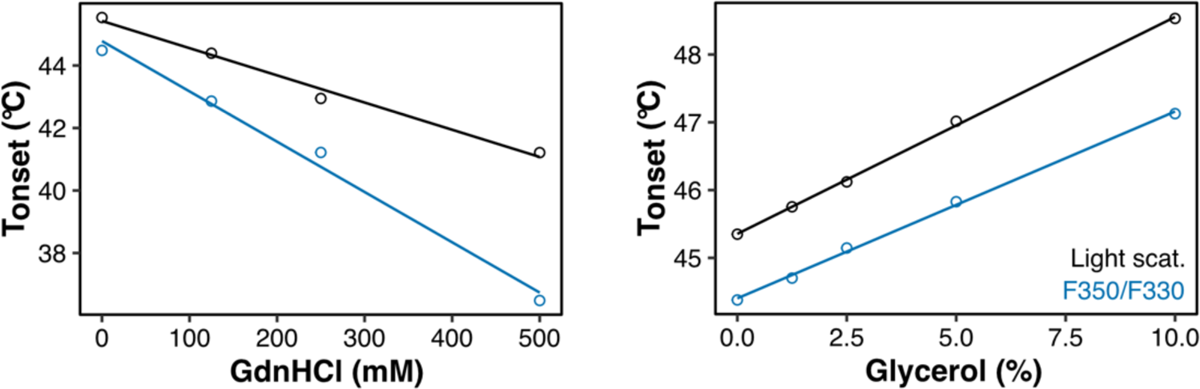
Chemical (de)stabilization of the structural stability alters the Ded1p assembly temperature. Change in T_onset_ for F350/F330 and light scattering signal for 4 μM GFP-labelled *S. cerevisiae* Ded1p in the presence of different concentrations of guanidine hydrochloride (GdnHCl, left) and glycerol (right) during a representative nanoDSF experiment. A trendline is shown as a guide (black for light scattering, blue for F350/F330).

### Surface-exposed residues in the helicase domain contribute to thermo-adaptation

Our data demonstrate that the temperature-induced assembly of *Sc* Ded1p is determined by the global stability of the helicase domain. Accordingly, the Ded1p orthologs exhibit distinct and different structural stabilities that correlate with their assembly temperature (**Figure 3A**). We hypothesized that (de)stabilizing point mutations within the helicase domain drive the adaptation of Ded1p assembly across fungal evolution. To identify amino acid substitutions determining the stability of the helicase domain of Ded1p, we calculated the evolutionary rate at every position in the helicase domain across the orthologous Ded1p sequences from our phylogenetic analysis (**Supplemental Figure** 1) and mapped these scores onto the predicted structure of *Sc* Ded1p (Jumper et al., 2021; Varadi et al., 2022). The central region of the predicted Ded1p structure, which represents the catalytic core of Ded1p and includes known ATP- and RNA-binding motifs (Linder and Jankowsky, 2011; Sengoku et al., 2006; Sharma and Jankowsky, 2014; Song and Ji, 2019), is highly conserved (**Figure 5A**). By contrast, the amino acid sequence at the surface of the helicase domain of Ded1p is evolutionary diverged (**Figure 5A**). We predicted that the less evolutionary constrained residues forming this surface could be altered during evolution to adapt the structural stability of the helicase domain.

**Figure 5.**
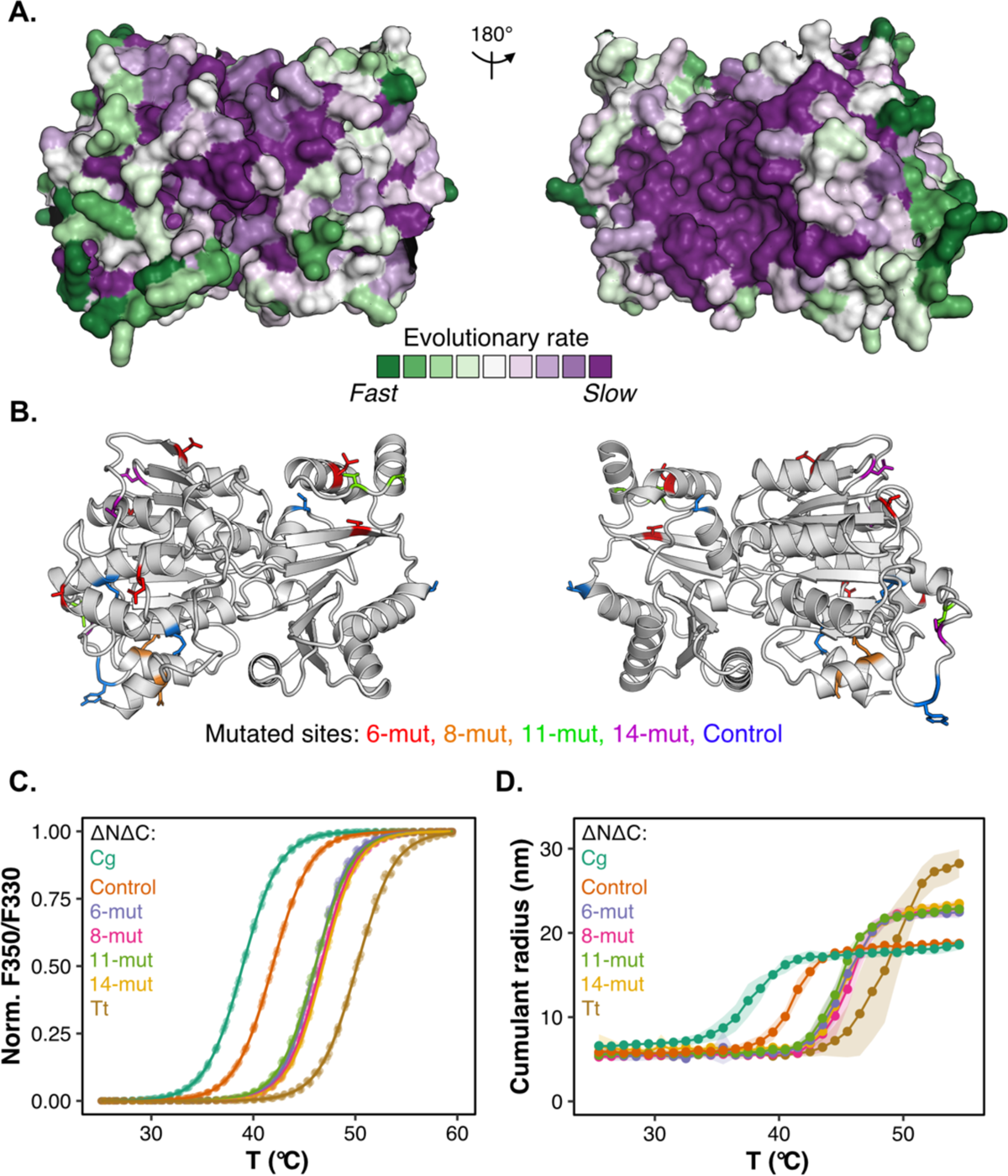
Adapting the assembly temperature of Ded1p by altering the structural stability of the helicase domain. **A.** AlphaFold structure of the helicase domain of *S. cerevisiae* Ded1p (Robinson, 2022; Varadi et al., 2022), in which surface colors represent differences in the evolutionary rate per residue. **B.** AlphaFold structure of the helicase domain of *C. globosum* Ded1p (Robinson, 2022; Varadi et al., 2022), highlighting residues that were predicted to be either “thermo-adaptive” (6-mut, 8-mut, 11-mut and 14-mut) or “non-thermo-adaptive” (Control) and mutated. **C.** Change in normalized F350/F330 as a function of temperature for 8 μM of the GFP-labelled helicase domains of *C. globosum* (C.g.) and *T. terrestris* (Tt) and variants (6-mut, 8-mut, 11-mut, 14-mut and Control). Mean (points), SD (light ribbon), n = 10. **D.** Change in cumulant radius (r_H_) as a function of temperature for 8 μM of the GFP-labelled helicase domains of *C. globosum* (C.g.) and *T. terrestris* (Tt) and variants (6-mut, 8-mut, 11-mut and Mock). Mean (points), sd (light ribbon), n = 4-5

To test this idea, we set out to increase the structural stability of the helicase domain of *Cg* Ded1p (mesophilic) towards that of *Tt* Ded1p (thermophilic). While these two species belong to the same fungal class and are relatively closely related (van Noort et al., 2013), the difference in structural stability of the full-length proteins is reflected by temperature transitions that are 13.2°C apart (**Figure 3A**). The amino acid sequence of the helicase domain of *Cg* and *Tt* Dedp1 differs in twenty positions (**Supplemental Figure 4A**). We compared these twenty positions to corresponding sites in Ded1p from other Sordariomycetes species and created a “ranking” of amino acid candidate substitutions (**Figure 1A, Supplemental Figure 4A**). We anticipated that if an amino acid residue is exclusively found in either thermo- or mesophilic Sordariomycetes orthologs, the substitution is more likely to be thermo-adaptive than if that residue occurs in both thermo- and mesophilic orthologs. Following this rationale, we designed four “thermo-adaptive” variants in which we sequentially reduced the number of substitutions between *Tt* and *Cg* Ded1p. In these variants, we replaced either fourteen (14-mut), eleven (11-mut), eight (8-mut) or six (6-mut) amino acid residues of *Cg* Ded1p with the corresponding residues from *Tt* Ded1p (**Figure 5B** and **Supplemental Figure 4A-B**). As a control, we designed a “non-thermo-adaptive” variant (Control) containing six substitutions that naturally occurred in the evolution of *Cg* and *Tt* Ded1p orthologs but are neither exclusively found in other closely related meso- and thermophilic Ded1p orthologs (**Figure 5B, Supplemental Figure 4A**). Most of the thermo- and non-thermo-adaptive candidate sites identified in the two Sordariomycetes orthologs were located on the more variable helicase domain surface (**Figure 5A-B**).

Next, we expressed and purified the helicase domains of *Cg* and *Tt* as well as *Cg*-based engineered helicase variants and determined the fluorescence temperature profiles. The T_onset_ of the temperature-transitions of the isolated helicase domains of *Cg* and *Tt* differed by ∼11.2°C (**Figure 5C, Supplemental Figure 4C**), which is comparable to the difference observed with the full-length proteins (13.2°C, **Figure 3A**). For the six “non-thermo-adaptive” substitutions (Control, T_onset_ = 38.2°C ±0.2), the apparent T_onset_ of the structural change of the *Cg* helicase domain (T_onset_ = 35.3°C ±0.2) increased by 2.9°C, suggesting that (some of) these amino acid substitutions marginally increase Ded1p’s structural stability. Introducing fourteen substitutions that we considered “thermo-adaptive” (14-mut), increased the apparent T_onset_ of the engineered *Cg* helicase domain by 8.1°C (T_onset_ = 43.4 ±0.3°C, **Figure 5C, Supplemental Figure 4C**), indicating that these residues shift the T_onset_ by more than half of the observed difference between *Cg* and *Tt* Ded1p. In comparison to 14-mut, titrating the number of substitutions down to eleven (11-mut, T_onset_ = 42.5°C ±0.2°C), eight (8-mut, T_onset_ = 43.2°C ±0.3°C) or six substitutions (T_onset_ = 42.8 ±0.3°C) did not alter the T_onset_ much (**Figure 5C, Supplemental Figure 4C**). This suggests that the key stability-determining residues are in the set of six “thermo-adaptive” amino acid substitutions (6-mut).

Next, we analyzed the assembly temperatures for the respective variants (**Figure 5D**). In agreement with our fluorescence data, residues which we considered to be non-thermoadaptive increased the assembly onset temperature by 2.9°C (*Cg* 34.4°C ±0.9°C; Control 37.3°C ±0.6°C). In contrast, the thermo-adaptive residues increased the assembly transition temperature by ∼8°C (6-mut (42.2°C ±0.8°C), 8-mut (42.6°C ±1.3°C), 11-mut (41.8°C ±0.4°C) and 14-mut (43.2°C ±0.4°C)) (**Figure 5D**). In summary, we identified a set of six residues which we consider “thermo-adaptive”. These residues appear to be sufficient to increase the stability of the helicase domain as well as the assembly temperature of Ded1p. This suggests that an evolutionary route to adapting the assembly temperature of Ded1p is by changing the temperature stability of the helicase domain.

### IDR-helicase domain chimeras reveal interactions between the structured domain and the N-terminal IDR

Recently, IDRs have been discussed as drivers, as well as modulators of condensate assembly. Our data indicate that the heat-induced assembly of Ded1p is primarily determined by the structural stability of the helicase domain (**Figure 5D**). However, our analysis also revealed that *Sc* Ded1p deletion variants lacking the N-terminal IDRs are structurally destabilized compared to wildtype *Sc* Ded1p (**Figure 3B-C**).

To characterize the effect of the IDRs on the structural stability and assembly temperature of Ded1p, we designed chimeric proteins in which we substituted either the N-terminal, C-terminal or both IDRs from mesophilic *Sc* Ded1p with those of thermophilic *Tt* Ded1p. To name the chimeric proteins, we use a three-letter code in which the order of letters refers to the domain architecture of Ded1p (N-terminal IDR, helicase domain and C-terminal IDR respectively) and the letter to the species from which the domain was taken (‘M’ for mesophilic, ‘T’ thermophilic) (**Figure 6A, Supplemental Figure 5A**). Consistent with removal of the C-terminal IDR (**Figure 3B-C**), substituting the mesophilic C-terminal IDR with the corresponding domain of thermophilic Ded1p (“MMT”) did neither substantially alter the structural stability nor the assembly temperature of Ded1p (**Figure 6B-C**), suggesting that the C-terminal IDR does not affect the stability of Ded1p. Instead, substituting the mesophilic N-terminal IDR with the thermophilic N-terminal IDR in TMM (T_onset_ light scattering = 40.2°C ± 0.2°C, T_onset_ F350/F330 = 38.9°C ±0.3°C) and TMT (T_onset_ light scattering = 39.6°C ± 0.4°C, T_onset_ F350/F330 = 40.0°C ±0.1°C) decreased the structural stability and assembly temperature of mesophilic Ded1p (T_onset_ light scattering = 44.2°C ± 0.1°C, T_onset_ F350/F330 = 42.7°C ±0.2°C) by ∼4°C (**Figure 6B-C**). The effect of this domain replacement is comparable to the removal of the N-terminal IDR as seen previously (**Figure 3B-C**) and demonstrates that the thermophilic N-terminal IDR cannot compensate for the mesophilic N-terminal IDR.

**Figure 6.**
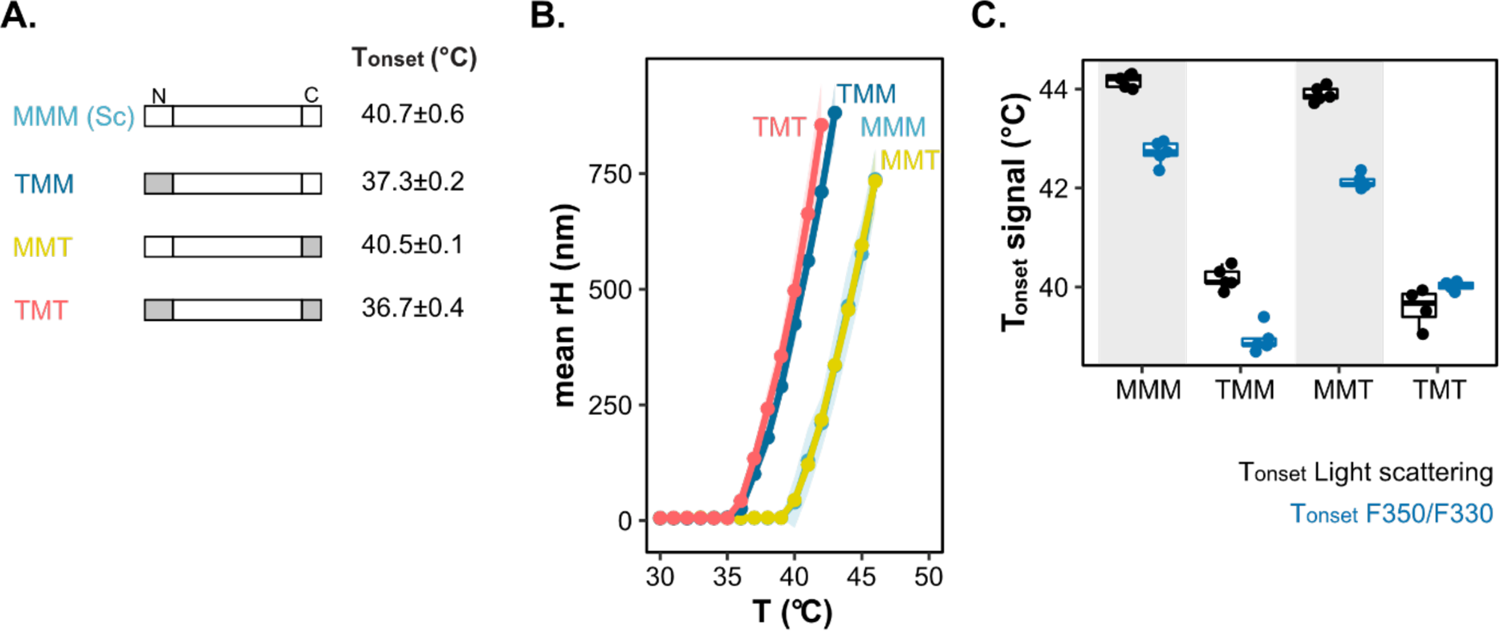
The helicase domain and IDRs together determine thermo-adaptation of Ded1p assembly temperature. **A.** Domain architecture of Ded1p from the mesophile *S. cerevisiae* (MMM (Sc)) and chimeric proteins in which either the N-terminal IDR (TMM), C-terminal IDR (MMT) or both IDRs (TMT) were exchanged. **B.** Change in hydrodynamic radius (r_H_) as a function of temperature for GFP-labelled Ded1p from *S. cerevisiae* (meso) chimeric proteins (see panel A) using DLS. Mean (points), sd (light ribbon), n = 3-4. **C.** Apparent T_onset_ of F350/F330 (blue) and back-reflection light scattering (black) signal for 5 μM of GFP-labelled Ded1p orthologs from *S. cerevisiae* (Meso) and chimeric proteins (see panel A) (n = 4-5).

To test if the N-terminal IDR from a species more closely related to *Sc* than *Tt,* could compensate for the N-terminal IDR of *Sc* Ded1p, we designed chimeric proteins of *Sk* (‘C’ for cold-adapted) and *Sc* Ded1p (‘M’ for mesophilic) (**Supplemental Figure 5B**). Again, exchanging the C-terminal IDR from *Sc* Ded1p with that of *Sk* Ded1p (MMC) did neither affect the structural stability nor the assembly temperature of *Sc* Ded1p (**Supplemental Figure 5C**). Substituting the N-terminal IDR in addition to the C-terminal IDR (CMC) reduced the assembly temperature of *Sc* Ded1p by ∼1.6°C and the structural stability by ∼2.5°C (**Supplemental Figure 5C**). This demonstrates that the structural stability and the assembly temperature of Ded1p is tuned by the N-terminal IDR (**Figure 6, Supplemental Figure 5**), which cannot be restored by substituting the N-terminal IDR with an orthologous N-terminal IDR, irrespective of the sequence divergence. This suggests that specific complementary stabilizing interactions exists between the disordered N-terminal region and the folded helicase domain of *Sc* Ded1p.

### The Ded1p N- and C-terminal IDRs dynamically interact with the helicase core

Our data so far suggest that the N-terminal IDR of *Sc* Ded1p interacts with the helicase domain. To test for these interactions experimentally, we applied NMR spectroscopic methods. To this end, we labeled the N-terminal IDR of *Sc* Ded1p with NMR active nuclei and recorded proton-nitrogen based NMR spectra to visualize H-N atom pairs (e.g., in a backbone amide) in the protein. The NMR spectra (**Figure 7A, black resonances**) confirmed that the N-terminal IDR does not adopt a stable tertiary structure, as the proton chemical shift dispersion was limited to a small region around 8.0 ppm. Addition of the NMR invisible *Sc* helicase domain (RecA1-RecA2) to the N-terminal IDR results in chemical shift perturbations (CSPs) and line broadening for a set of the resonances of the N-terminal IDR, revealing a direct and specific interaction between the two regions of the enzyme (**Supplemental Figure 6A).** We next narrowed down the region of the helicase core that interacts with the N-terminal IDR by performing NMR titration experiments with the isolated RecA1 (**Supplemental Figure 6B**) and Rec2 (**Figure 7A**) subdomains. This revealed that the N-terminal IDR-helicase core interaction is mediated by the RecA2 domain (**Figure 7A**) and implies that the RecA2 domain binds directly to the N-terminal IDR at a specific site. Next, we labeled the RecA2 domain with NMR active nuclei to verify the N-terminal IDR:RecA2 domain interactions in the other direction. 2-dimensional H-N spectra of the RecA2 domain in the absence and presence of the N-terminal IDR (**Figure 7B**, black and orange resonances, respectively), confirmed that a set of RecA2 resonances were altered upon the addition of the unlabeled N-terminal IDR. Interestingly, the extent of the CSPs was reduced at elevated temperatures (**Figure 7B**, compare 5°C with 25°C), demonstrating that the interactions between the N-terminal IDR and RecA2 domain weaken upon temperature increase. To identify the residues in the N-terminal IDR and RecA2 domains that mediate the interaction, we assigned the NMR resonances of both Ded1p fragments. This revealed a short sequence stretch between residues 20 and 30 within the N-terminal IDR (**Figure 7C**) that interacts directly with a surface exposed region in the RecA2 domain that includes and surrounds the helix between residues 380 and 397 (**Figure 7D**). Taken together, our NMR data revealed a short sequence within the N-terminal IDR that docks onto a surface patch of the RecA2 domain.

**Figure 7.**
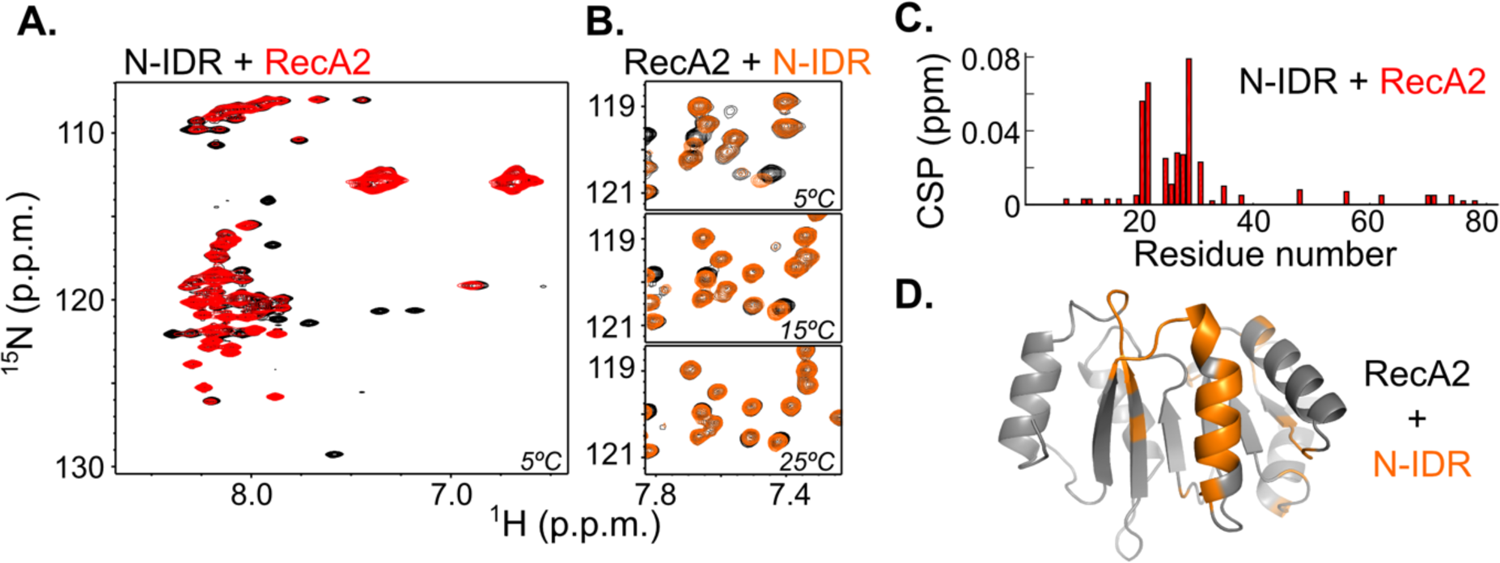
The IDRs interact with the helicase domain. **A.** ^1^H-^15^N NMR spectrum of the N-terminal IDR of Ded1p in the absence (black) and presence (red) of the Ded1p RecA2 domain. Several Ded1p resonances experience chemical shift perturbations (CSPs) and broaden upon interaction with the second RecA2 domain, indicating a specific binding event. **B**. ^1^H-^15^N NMR spectrum of the Ded1p RecA2 domain in the absence (black) and presence (orange) of the Ded1p N-terminal IDR. The extend of the chemical shift perturbations significantly decrease with increasing temperature, indicating that the interaction is weaker at higher temperatures. **C.** CSPs from panel A plotted against the sequence of the N-terminal IDR. Residues 20 to 30 in the N-terminal IDR interact directly with the RecA2 domain, as resonances of those residues experience the largest CSPs. **D.** CSPs from panel C plotted on a homology model of the Ded1p RecA2 domain. A specific surface patch around residues 380-397 on the Ded1p RecA2 domain (colored orange) is responsible for the interaction with the Ded1p IDR.

Next, we performed NMR titration experiments to reveal if and how the C-terminal IDR interacts with the helicase core. Upon addition of the helicase core to the C-terminal IDR the C-terminal IDR resonances broaden significantly, which reveals that the C-terminal IDR directly interacts with the helicase core domain (**Supplemental Figure 6C**). This interaction is mediated by the RecA1 domain, as addition of this domain also results in line broadening of the C-terminal IDR resonances, whereas addition of the RecA2 domain has no influence on the C-terminal IDR resonances. However, in this case, we were unable to localize the interaction to specific sites, as all resonances of C-terminal IDR domain broaden beyond detection upon complex formation. We attribute this this effect to the visible condensation that took place in the NMR tube at the concentrations required for the NMR experiments. In summary, our NMR titration data reveals a situation where the RecA1 domain in the helicase core interacts with the C-terminal IDR, whereas the RecA2 domain specifically binds to the N-terminal IDR (**Supplemental Figure 6D)**.

### The IDRs mediate micron-sized condensate assembly of Ded1p

Our NMR data show that the N-terminal and C-terminal IDR are folded back onto the helicase domain, and the interaction between the N-terminal IDR and the helicase domain weakens with increasing temperatures (**Figure 7B**). We speculate that upon exposure to heat, the IDRs become available to establish intermolecular interactions that drive condensation. Consistent with this reasoning, DLS data reveal that full-length Ded1p assembles into micron-sized condensates upon heating, whereas variants in which the N-terminal, C-terminal or both IDRs were removed assemble into particles that are at least one order of magnitude smaller (**8A-B**). These data suggest that the IDRs provide protein-protein interactions that provide additional valences for growth into condensates.

To identify residues that could be important for establishing protein-protein interactions for condensation, we returned to our phylogenetic analysis. While tryptophan residues are typically excluded from IDRs, we found that the tryptophan residues (or aromatic residues in general) in the C-terminal IDR of *Sc* Ded1p are highly conserved (**Supplemental Figure 7**).

To test whether the C-terminal tryptophan residues influence the heat-induced assembly of *Sc* Ded1p, we substituted five tryptophan residues in the C-terminal IDR with alanine residues, giving rise to Ded1p-5WA. The T_onset_ for the structural change of Ded1p-5WA was comparable to that of wildtype Ded1p (**Figure 9A**). However, the assembly temperature of Ded1p-5WA was increased compared to wildtype Ded1p (**Figure 9B**), suggesting that the tryptophan residues within the C-terminal IDR contribute to protein-protein interactions for Ded1p condensate assembly. In summary, these data indicate that heat-induced assembly of Ded1p into condensates is regulated by a complex interplay of the IDRs and the structured domain, which is subject to evolutionary tuning.

## Discussion

In this paper, we describe a molecular mechanism for detecting and responding to temperature changes by Ded1p, an essential translation factor that assembles into condensates regulating the HSR (Iserman et al., 2020). Our data suggest that heat triggers a conformational change in the folded helicase domain of Ded1p, which promotes assembly into nanometer-sized particles. This is accompanied by changes in the conformational ensemble of the flanking IDRs, which facilitates assembly into micron-sized condensates. Ded1p assembly into condensates is conserved among various fungi and the assembly temperature is adapted to the species’ growth temperature, suggesting that Ded1p’s function in the HSR is conserved and adjusted to a species’ temperature niche. Given that many other translation factors co-assemble into condensates upon exposure to heat and are composed of both folded and disordered domains, our data suggest a general model for temperature-induced protein assembly and thermo-adaptation of the translational HSR.

The helicase domain of Ded1p consists of two globular RecA domains that each adopt a Rossmann fold. The secondary structure of these folds is largely maintained upon temperature increase (Iserman et al., 2020), suggesting that the helicase domain does not undergo extensive denaturation. Here, using nanoDSF, a method that is sensitive to tertiary structure changes, we identified a heat-induced structural change in the helicase domain that coincides with assembly (**Figure 3A-B**). This Ded1p conformational change has also been observed in budding yeast cell extracts using limited proteolysis and mass spectrometry (Cappelletti et al., 2021; Leuenberger et al., 2017). These studies detected regions in the helicase domain of Ded1p that become increasingly exposed to proteolysis upon heating. Two of these regions, amino acids 136-158 (Leuenberger et al., 2017) and amino acids 207-222 (Cappelletti et al., 2021), reside in the RecA1 and are in proximity of a single tryptophan residue that reports on the structural changes that we detect in our nanoDSF experiments. The heat-induced change in tryptophan fluorescence detected with nanoDSF could thus indicate a local displacement of these two regions. We speculate that this heat-induced structural change exposes hidden sites in the helicase domain that provide valences for assembly (Ruff et al., 2022). The chemical nature of these valences is unclear and remains to be investigated.

We previously showed that Ded1p assembly is linked to regulating the HSR (Iserman et al., 2020). Given the conservation and thermal adaptation of the HSR, we tested whether the mechanism of Ded1p assembly is also adapted to the thermal niche of a given species. Using *in vitro* condensation experiments with various fungal Ded1p orthologs, we were able to show that the structural stability of Ded1p correlates with a species’ growth temperature (**Table 1**).

In fact, all tested orthologs contain a single conserved tryptophan residue in their helicase domain (253W in *Sc* Ded1p) and exhibit similar spectroscopic shifts that presumably represents a similar tertiary structure change (**Figure 3A** and **Supplemental Figure 3C**). Importantly, the thermostability of *Sc* Ded1p is smaller compared to the rest of the proteome, and it changes its tertiary structure well below the lethal temperature of the organism (Iserman et al., 2020; Wallace et al., 2015). This suggests that evolutionary pressures are acting to finetune the structural stability of Ded1p throughout fungal evolution, which is most likely linked to Dep1p’s role in regulating the translational HSR.

Our data suggest that just a few amino acid substitutions can render the helicase domain of Ded1p more stable. Substituting six residues in the helicase domain of mesophilic Ded1p with the corresponding residues from thermophilic Ded1p was sufficient to increase the structural stability by more than 7°C (**Figure 5C, Supplemental Figure 4C**). We did not observe similar amino acids changes in distantly related orthologs (**Supplemental Figure 4B**), suggesting that this may be a unique evolutionary route for thermo-adapting Ded1p. This is consistent with previous work, where thermo-adaptive amino acid changes were mapped to various parts of lactate dehydrogenase, an enzyme that shares the Rossmann fold. Importantly, few of the different thermo-adaptive amino acid changes found in LDH orthologs affect the stability of the same structural or functional elements (Fields et al., 2015), suggesting that there are multiple paths for adapting structural protein stabilities. Our limited set of variants already indicates that the stability of Ded1p can readily be altered by evolution. The fact that *Sc* Ded1p has not become more stable over long evolutionary time scales, suggests that a higher Ded1p structural stability is unfavorable, most likely because it is tightly coupled to a physiologically important temperature threshold for regulating the HSR.

Most DEAD-box helicases contain accessory domains in addition to their helicase domains. The accessory domains of Ded1p, the N-terminal and C-terminal IDRs, were shown previously to be required for Ded1p’s role in translation initiation in growing cells (Floor et al., 2016). The N-terminal IDR is prion-like in composition with a high content of glycine, arginine, and polar residues, such as serine and asparagine. Prion-like domains have been implicated in promoting assembly via amyloid and non-amyloid-like interactions (Franzmann and Alberti, 2019b), but other roles have been proposed as well (Vijayakumar et al., 2019). Our NMR data show that the N-terminal prion-like IDR folds back onto the RecA2 domain (**Figure 7A-B**). This interaction stabilizes the helicase domain at physiological temperatures and removal or exchange of the N-terminal IDR with an IDR from related species results in reduced Ded1p stability (**Figure 3B and Figure 6C**). In addition to its stabilizing function, the interaction between the N-terminal IDR and the helicase domain, as well as its interactions with other proteins are likely relevant in growing cells, as N-terminal truncation variants confer slow growth phenotypes in yeast (Floor et al., 2016). We have been unable to map the interaction sites of the C-terminal IDR and the helicase domain in our NMR experiments due to the propensity of this IDR to assemble into condensates. However, Alphafold predicts a region in the C-terminal IDR that contacts the helicase domain (Robinson, 2022; Varadi et al., 2022) (**Supplemental Figure 8**), albeit with low confidence.

This region contains a highly conserved motif (RDYR), which is a potential candidate for interacting with the helicase domain. While the C-terminal IDR does not seem to affect the thermostability of Ded1p much (**Figure 3B**), future work will focus on investigating the role of this motif in regulating Ded1p’s function. Interestingly, the Ded1p Alphafold model also predicts with low confidence, an interaction between the N-terminal IDR and the second RecA2 domain. The computationally calculated binding sites on the RecA2 and N-IDR are surprisingly close and in full agreement with our experimental data on this interaction (**Figure 7D)**.

Both the N- and C-terminal IDRs are required for Ded1p to assemble into micron-sized condensates (**Figure 8A-B**), and we speculate that heat-induced conformational changes (such as weakening of the interaction between the N-terminal IDR and the helicase domain) expose valences in these IDRs that facilitate higher-order assembly. Such valences can be organized in sticker and spacer regions, in which adhesive residues are spatially segregated by less-adhesive linker regions (Harmon et al., 2017). Examples of sticker residues in Ded1p are the aromatic residues, which are regularly spaced throughout the C-terminal IDR. Indeed, substituting the C-terminal tryptophan residues with alanine residues results in a reduction in the propensity of Ded1p to assemble, as evidenced by an increased assembly temperature (**Figure 9B**), without affecting the onset temperature of the structural change much (**Figure 9A**). Given that the N- and C-terminal IDRs contain binding sites for various components of the translation pre-initiation complex (Gao et al., 2016; Gulay et al., 2020; Hilliker et al., 2011), temperature-driven assembly of Ded1p may be modulated by association with other translation initiation factors. It is also conceivable that these translation initiation factors promote additional heterotypic interactions for example with RNA that drive assembly into multicomponent condensates.

**Figure 8.**
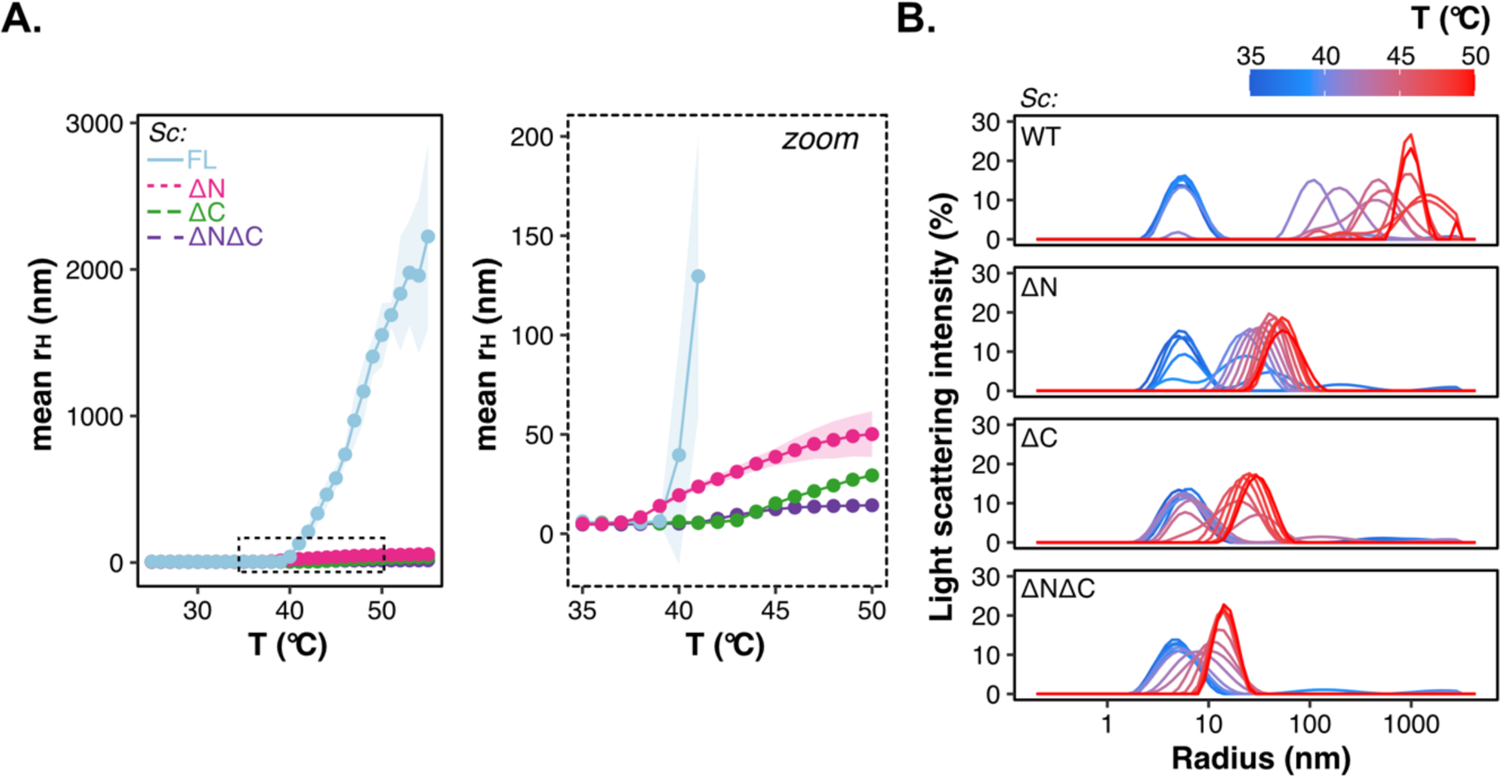
The IDRs provide valences for micron-sized assembly upon heating. **A.** Change in hydrodynamic radius (r_H_) as a function of temperature for GFP-labelled full length *S. cerevisiae* Ded1p (FL), Ded1p-ϕλN, -ϕλC and -ϕλNϕλC using DLS. Inset shows zoom at r_H_ < 200 nm. Mean (points), sd (light ribbon), n = 3-4. **B.** Representative DLS experiment showing the distribution of the light scattering intensities as a function of particle size at different temperatures for GFP-labelled full length *S. cerevisiae* Ded1p, Ded1p-ϕλN, -ϕλC and -ϕλNϕλC.

**Figure 9.**
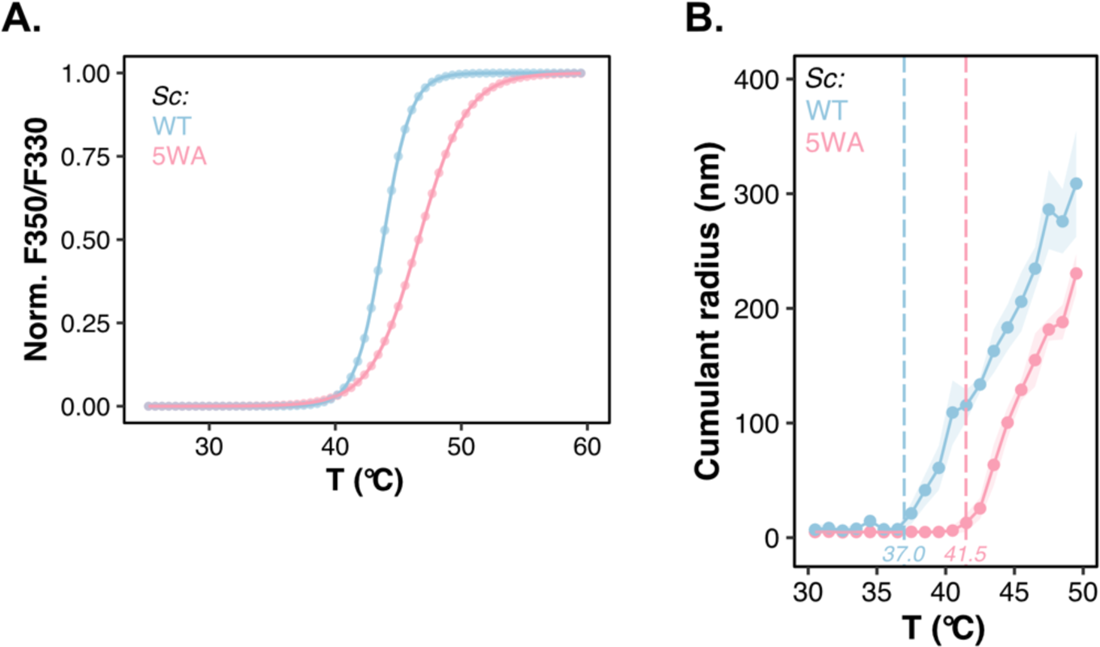
Conserved Trp residues in the C-terminal IDR influence the Ded1p assembly temperature. **A.** Change in normalized F350/F330 as a function of temperature for 10 μM unlabeled wildtype *S. cerevisiae* Ded1p (WT) and Ded1p-5WA (5WA) using nanoDSF. A trendline is shown as a guide. Mean (points), sd (light ribbon), n = 2-4. **B.** Change in hydrodynamic radius (r_H_) as a function of temperature for GFP-labelled *S. cerevisiae* Ded1p (WT) and Ded1p-5WA (5WA) using DLS. Mean (points), sd (light ribbon), n = 4.

**Figure 10.**
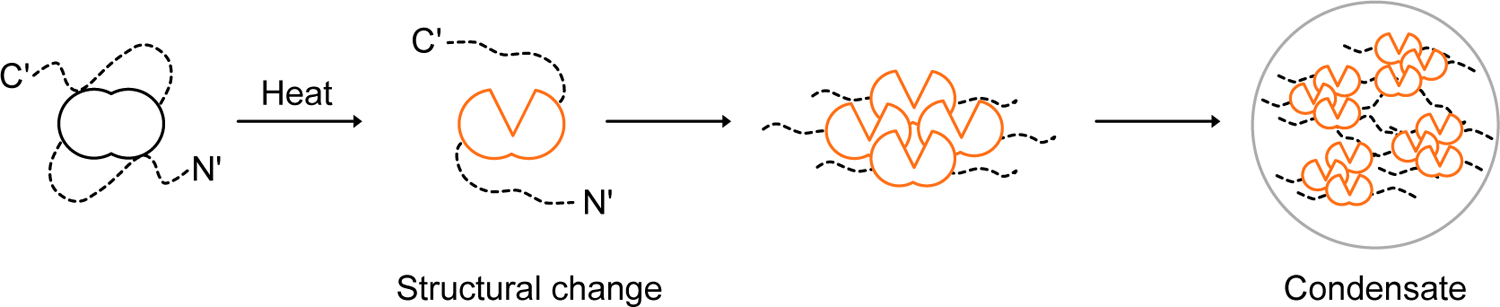
Model for heat-induced Ded1p assembly. Under non-stress conditions, Ded1p exists in a diffuse state, in which the structured helicase domain interacts with the N- and C-terminal IDRs. Upon exposure to heat, the structured helicase domain undergoes a change in tertiary structure, and this is sufficient to induce the assembly of nanometer-sized particles. Because of the heat-induced structural change, the IDRs are now available to promote intermolecular assemblies that promote assembly into larger, micron-sized particles. To adapt the assembly temperature of Ded1p to different temperatures, the structural stability of the helicase domain must co-evolve with the IDRs.

Previous plant work implicated prion-like IDRs as thermo-responsive devices that detect temperature changes by forming condensates (Jung et al., 2020; Wilkinson and Strader, 2020), but the molecular mechanisms underlying condensation have not yet been worked out. Our results are consistent with these studies but suggests that IDRs are not essential for heat-induced assembly of Ded1p (**Figure 8A-B**). Rather, condensate assembly by Ded1p requires a concerted action of the structured helicase domain and IDRs. This synergy between folded and disordered domains provides an explanation for the observed co-evolution of these domains in *Sc* Ded1p to set the assembly temperature (**Figure 6**). Notably, this coupling between the disordered and folded domains is reminiscent of reports focusing on other translation factors in budding yeast. For example, heat-induced assembly of the poly(A)binding protein (Pab1p) is determined by the structured RRM domain and adjusted to physiological temperatures by a disordered proline-rich IDR (Riback et al., 2017). Similar findings were made for the functionally related protein Pub1p (Kroschwald et al., 2018). Furthermore, pH-induced assembly of Sup35 requires interactions between a the structured GTPase domain and residues in a prion-like N-terminal IDR (Franzmann et al., 2018). We thus propose that the coupling of conformational changes in structured domains and IDRs constitutes a common mechanism for responding to temperature changes and the formation of condensates.

Heat-induced Ded1p assembly is reversible in yeast, yet reconstituted Ded1p condensates are not reversible by lowering the temperature (Iserman et al., 2020), suggesting that cellular components are necessary for Ded1p disassembly. Indeed, previous work in cells has shown that dissolution of heat-induced translation factor condensates requires molecular chaperones (Cherkasov et al., 2013; Kroschwald et al., 2018; Yoo et al., 2022). We speculate that the disassembly of Ded1p condensates *in vitro* requires the reversal of temperature-induced structural changes in the helicase domain by molecular chaperones.

In summary, we propose a model for temperature-controlled assembly of Ded1p that involves coupling of interdomain communication and conformational changes in the structured domain and disordered regions. Our work provides a molecular understanding for how organisms can respond to temperature changes and gives insights into the evolutionary routes that are available for organisms to adapt to new climates.

## Acknowledgements

We thank the following MPI-CBG facilities: Protein Expression Purification and Characterization, Light Microscopy, FACS and Scientific Computing. We thank the Alberti and Hyman lab members for comments on the manuscript and Dorothee Thiel support with insect cell culture. We are grateful to Rohit Pappu for insightful discussions on the mechanism of Ded1p condensation. We thank Nanotemper Technologies GmbH for providing early access to Prometheus Panta.

## Author contributions

CJ: Conceptualization, methodology, validation, formal analysis, investigation, writing original draft, writing review & editing, visualization, project administration, funding acquisition

TF: Conceptualization, methodology, validation, formal analysis, investigation, writing review & editing, supervision

JH: Methodology (NMR), validation, investigation JS: Methodology (NMR), validation, investigation

CL: Methodology, validation, investigation, visualization SW: Methodology, validation, supervision

ATP: Methodology, writing review & editing, supervision, funding acquisition

RS: Methodology (NMR), formal analysis, writing review & editing, visualization, supervision, funding acquisition

AAH: Conceptualization, writing review & editing, supervision, funding acquisition

SA: Conceptualization, writing original draft, writing review & editing, supervision, funding acquisition

## Materials and methods

### Phylogenetic analysis and proteomic comparison

The proteomes of fungi belonging to the Saccharomycotina clade were downloaded from Shen et al. (2018) (Shen et al., 2018) and Morgenstern et al. (2012) (via CSFG Genomes) (Morgenstern et al., 2012). In addition, all eukaryotic, non-redundant, and non-excluded proteomes were downloaded from Uniprot (UniProt Consortium, 2019). All sequences from all proteomes were blasted against all other proteomes using pOrthoMCL (Tabari and Su, 2017) to identify Ded1p orthologs. The reverse best hit was taken at a normalized edge width of;: 1.5, followed by sequence alignment. After manual cleaning, the alignment contained 630 sequences.

A phylogenetic tree was constructed using IQ-TREE (1.6.11) (Lawson Handley et al., 2019), in which the amino acid sequence of the human homolog DDX3X was used as an outgroup. To find the optimal transition matrix, selection was performed on 546 models. The best model was LG+F+R10; LG: general matrix (Le and Gascuel, 2008), F: empirical amino acid frequencies from the data, R10: FreeRate model which discretizes the gamma-like distribution into 10 rate categories (Soubrier et al., 2012; Yang, 1995). Node support was determined by bootstrap (UFBoot; 10,000 x) (Hoang et al., 2018).

Site specific evolutionary rates were calculated using IQTree’s (2.1.2) (Minh et al., 2020) Bayesian rate inference and the previously determined best fitting model LG+F+R10. The same 630 sequences were aligned with only sites in DED1_YEAST preserved. Site specific rate categories were used to color the Ded1p structure taken from AlphaFold (Robinson, 2022; Varadi et al., 2022). Rate categories 9 and 10 were combined. A sequence logo was plotted using the R (version 4.0) package ggseqlogo (version 0.1), in which the hight of the letters represents the frequency of occurrence at a given position in the alignment.

### Yeast culture and spotting assay

*S. cerevisiae, S. kudriavzevii* and *O. parapolymorpha* were grown to exponential phase in YPD medium at 30°C. The cell density of yeast cultures was assessed by measuring OD_600_ and cultures were adjusted to equal cell concentrations. The cells were diluted in fivefold serial dilutions with YPD in 96-well plates (conical bottom, Sarstedt). Diluted cultures were spotted onto YPD agar plates using a sterilized multi-blot 48 pin replicator. The plates were incubated for 24 h at the indicated temperatures and photographed with a standard digital camera.

### Protein construct design

#### S. cerevisiae Ded1p

The helicase domain boundaries of Ded1p from *S. cerevisiae* (**Table 2-3**) were determined by aligning the amino acid sequence of Ded1p to crystal structures of the human ortholog DDX3X, including PDB 2I4I, 4PXA and 5E7I (Floor et al., 2016; Högbom et al., 2007; (Epling et al., 2015). The amino acid sequence of Ded1p that aligned with the structured region of the DDX3X crystal structures (143-578) was defined here as the structured helicase domain of Ded1p (**Table 3**) and these amino acid coordinates were used to design of truncation variants and chimeric proteins in this paper (**Table 3-4)**. The N-terminal and C-terminal regions that flank the Ded1p helicase domain of are predicted to lack defined secondary structure (Buchan and Jones, 2019) and to be intrinsically disordered (Dosztányi et al., 2005).

**Table 2.** Strain and protein identifiers of Ded1p orthologs used in this study.

| Ded1p ortholog from: | Fungal strain identifier | Uniprot identifier |
| --- | --- | --- |
| *S. cerevisiae* | S288C | P06634 |
| *S. kudriavzevii* | IFO1802 | J4U4J8 |
| *O. parapolymorpha* | DL-1 | W1QIJ9 |
| *C. globosum* | CBS148.51 | Q2HBE7 |
| *T. terrestris* | NRRL8126 | G2QTC2 |

**Table 3.** Domain boundaries for the Ded1p orthologs used to create chimeric proteins.

|  | N-terminus | Helicase domain | C-terminus |
| --- | --- | --- | --- |
| Ded1p *S. cerevisiae* | 1-98 | 99-535 | 536-604 |
| Ded1p *S. kudriavzevii* | 1-102 | 103-539 | 540-609 |
| Ded1p *T. terrestris* | 1-143 | 144-582 | 583-666 |

**Table 4.** Design of Ded1p *S. cerevisiae* truncation variants

| Variant | Construct |
| --- | --- |
| Ded1p-ΔN (Sc) | Met-99-604 |
| Ded1p-ΔNΔC (Sc) | Met-99-535 |
| Ded1p-ΔC (Sc) | 1-535 |

The tryptophan variant Ded1p-5WA consists of the amino acid sequence of *S. cerevisiae* Ded1p with the following point mutations: W567A, W583A, W592A, W603A, W604A

#### Chimeric proteins

Chimeric proteins were designed by exchanging the N-terminus, helicase domain and/or C-terminus *from S. cerevisiae* Ded1p with the respective domains of Ded1p orthologs from *S. kudriavzevii* and *T. terrestris*. To determine the domain boundaries of Ded1p orthologs (Table 3), the orthologous amino acid sequences were aligned to the sequence of *S. cerevisiae* Ded1p.

#### C. globosum helicase domain variants

To study the thermo-adaptation of *Cg* Ded1p and *Tt* Ded1p, helicase domain variants were designed in which amino acids of *Cg* Ded1p were exchanged with the corresponding amino acid of *Tt* Ded1p (Table 5). These constructs contain the structured domain only and not the intrinsically disordered regions (Table 5). To identify candidate sites, the amino acid sequences of both *Cg* (mesophile) and *Tt* (thermophile) were aligned with orthologous sequences from other fungi from the Sordariomycetes class: *Thermothelomyces thermophilus* (thermophile, Uniprot ID G2Q8V8), *Chaetomium thermophilum* (thermophile, uniprot ID G0SG53) and *Neurospora crassa* (mesophile, uniprot ID Q9P6U9) (**Supplemental Figure 5B**). Increasing numbers of mutations were introduced into the helicase domain of *Cg* Ded1p (Table 5). As a control, six positions were mutated in which residues differ between *Cg* Ded1p and *Tt* Ded1p but are similar between *Cg* Ded1p and orthologs from other thermophilic Sordariomycetes (see above) and/or positions in which the residue in *N. crassa* Ded1p is similar to that of *Tt* Ded1p.

**Table 5.**
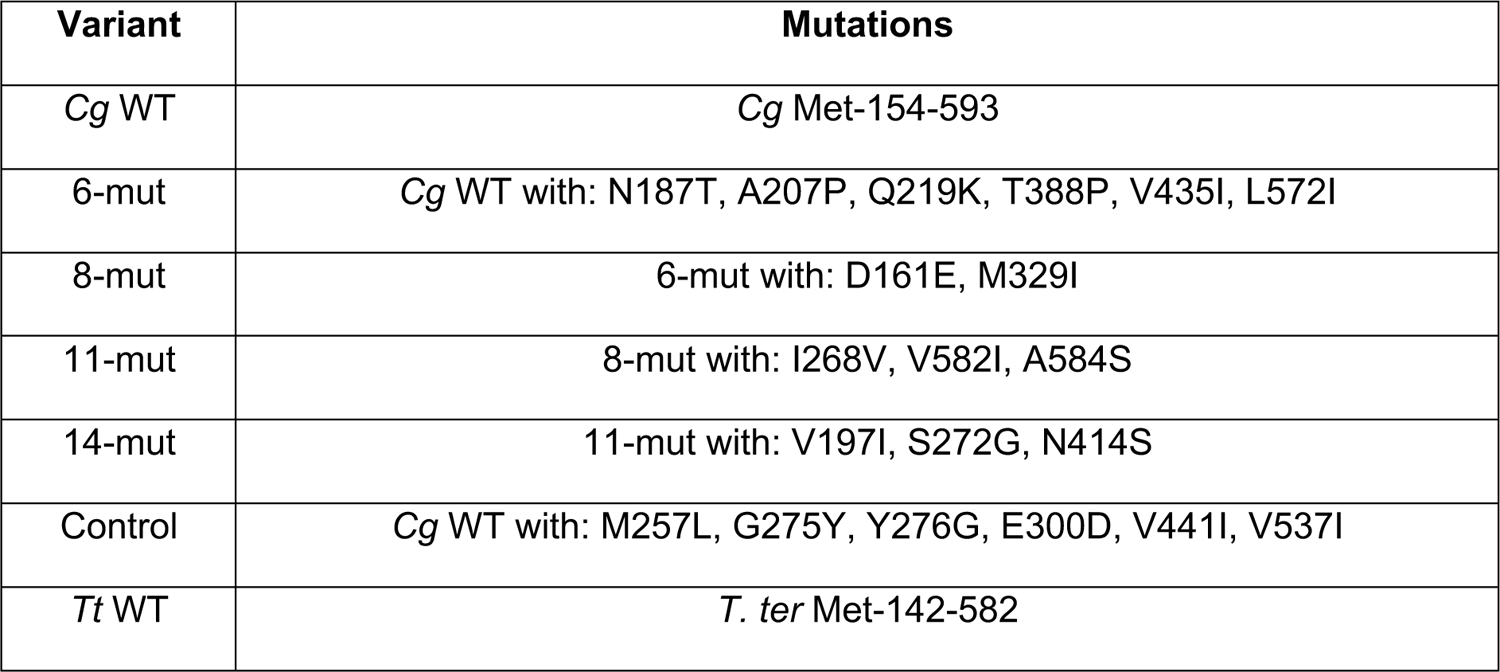
Overview of the mutations in the helicase domain of *Cg* Ded1.

| Variant | Mutations |
| --- | --- |
| *Cg* WT | *Cg* Met-154-593 |
| 6-mut | *Cg* WT with: N187T, A207P, Q219K, T388P, V435I, L572I |
| 8-mut | 6-mut with: D161E, M329I |
| 11-mut | 8-mut with: I268V, V582I, A584S |
| 14-mut | 11-mut with: V197I, S272G, N414S |
| Control | *Cg* WT with: M257L, G275Y, Y276G, E300D, V441I, V537I |
| *Tt* WT | *T. ter* Met-142-582 |

#### Cloning and mutagenesis

ORFs were codon optimized for *Spodoptera frugiperda* using IDT codon optimizer and ordered as gBlocks (IDT, Leuven, Belgium). gBlocks were cloned into pUC57-Kanamycin plasmids (Genscript, Rijswijk, Netherlands) by restriction/digestion (NEB, Frankfurt am Main, Germany) and quick ligation (NEB, Frankfurt am Main, Germany). Before ligation, the pUC57-Kanamycin were also digested by restriction enzymes (NEB, Frankfurt am Main, Germany) and gel extracted using the QIAprep Gel Extraction kit (QIAGEN, Hilden, Germany). The sequence was verified by colony PCR using standard primers (Sigma-Aldrich, Taufkirchen, Germany) and Taq polymerase (In house, MPI-CBG). *DED1* Trp-to-Ala point mutations were introduced using the Q5 site-directed mutagenesis kit (NEB, Frankfurt am Main, Germany) with primers carrying the sequence variation and verified by sequencing. For virus production, the genes were subcloned into pOCC shuttle vectors using the restriction sites AscI and NotI (NEB, Frankfurt am Main, Germany) (Lemaitre et al., 2019) and quick ligation (NEB, Frankfurt am Main, Germany). All plasmids were verified by sequencing.

### Bacterial transformation and plasmid extraction

Chemically competent *E. coli* DH5α were thawed and incubated with 10 μL ligation volume on ice. The cells were heat shocked at 42°C for 1 min, followed by 5 min incubation on ice. After 1 h recovery at 37°C at 700 rpm in lysogeny broth (LB) medium (1% peptone, 0.5% yeast extract, 1% NaCl), the bacteria were streaked on plates with Kanamycin (pUC57 plasmids) or Ampicillin (pOCC plasmids) and incubated at 37°C overnight. Clones were inoculated in 5 mL LB medium with antibiotics. After overnight culture at 37°C at 220 rpm, plasmids were extracted using the QIAprep Spin Miniprep Kit (QIAGEN, Hilden, Germany).

### Protein purification

Recombinant MBP-3C-DED1-monoGFP-3C-6xHis (Ded1p orthologs and variants) and MBP-3C-DED1-3C-6xHis (*S. cerevisiae* Ded1p, Ded1p-5WA) were expressed for 72 h in Sf9 insect cells using a baculovirus expression system (Lemaitre et al., 2019). Cells were lyzed at 10-15,000 PSI on ice using an LM20 Microfluidizer (Microfluidics, Westwood, USA) in 50 mM Tris/HCl pH 8.0, 1 M KCl, 2 mM EDTA, 1 mM DTT, Benzonase (MPI-CBG) and protease inhibitor (Merck, Darmstadt, Germany). The soluble fraction of the lysate was collected after centrifugation for 45 min at 20,000 rpm using a Ti-45 rotor or at 25,000 rpm using a JA25.50 rotor (Beckman Coulter, Krefeld, Germany) at 4°C. The supernatant of the cell lysate was filtered through a 0.22 μm filter (Corning, Kaiserslautern, Germany) and incubated with amylose resin (NEB, Frankfurt am Main, Germany) for 1 hour at 4°C. After washing on batch and on column, the protein was eluted with washing buffer (50 mM Tris/HCl pH 8.0, 1 M KCl, 2mM EDTA and 1 mM DTT) containing 20 mM maltose. His and MBP tags were cleaved off with His-tagged PreScission protease (3C, in-house MPI-CBG) at 4°C overnight. After concentrating the samples with 30 K Vivaspin centrifugal filters (Vivaproducts, Littleton, USA), the samples were applied to a HiLoad Superdex 200 pg 16/600, Superdex 200 pg 26/600 or Superdex 200 pg 10/30 (Cytiva, Marlborough, USA) equilibrated with washing buffer. Gelfiltration was carried out with an ÄKTA pure 25 (GE Life Sciences, Germany) at room temperature. Fractions were analyzed by SDS-PAGE, pooled and the protein concentrated to 30-120 μM using 30 K Vivaspin centrifugal filters (Vivaproducts, Littleton, USA). Aliquots were flash-frozen and stored at −80°C. For experiments, proteins were thawed and spun at 20,000 xg for 5 min at 20°C (Eppendorf microcentrifuge, Germany). The protein concentration was determined by measuring the absorbance at λ = 280 nm and/or λ = 488 nm (for GFP-labelled proteins) using a NP-80 UV/VIS nanospectrophotometer (Implen, Munich, Germany). Purified proteins were analyzed in 12.5% SDS-PAGE gels. Gels were stained with InstantBlue^TM^ (Abcam, Cambridge, UK) or with Coomassie G-250 (Lawrence and Besir, 2009).

### Dynamic light scattering (DLS)

Dynamic light scattering of 4 μM GFP-labelled purified protein was measured with a Zetasizer Nano ZSP (Malvern Panalytical Ltd., Malvern, UK) using 173° backward scattering angle. Proteins were tested in 19 μl reactions containing 80 mM phosphate buffer pH 7.4, 200 mM KCl and 0.4 mM TCEP. Measurements were recorded with the Ultra-Low volume quartz Zen2112 cuvette (Hellma, Jena, Germany) with lid to prevent evaporation. The samples were temperature-equilibrated at room temperature for ∼15 min. Eight five second autocorrelations were recorded, averaged and analyzed using the manufacturers software. For temperature experiments, the temperature was increased from 20°C to 60°C with 1°C increments and 30 s equilibration time at each temperature. For temperature experiments, the mean hydrodynamic radii were plotted as a function of temperature, typically between 30°C:−σ T:−σ 55°C and mean the r_H_ < 500 nm to visualize variant-specific differences in the assembly temperature (T_onset_). The assembly temperature (T_onset_) was derived from fitting the data to equation 1,

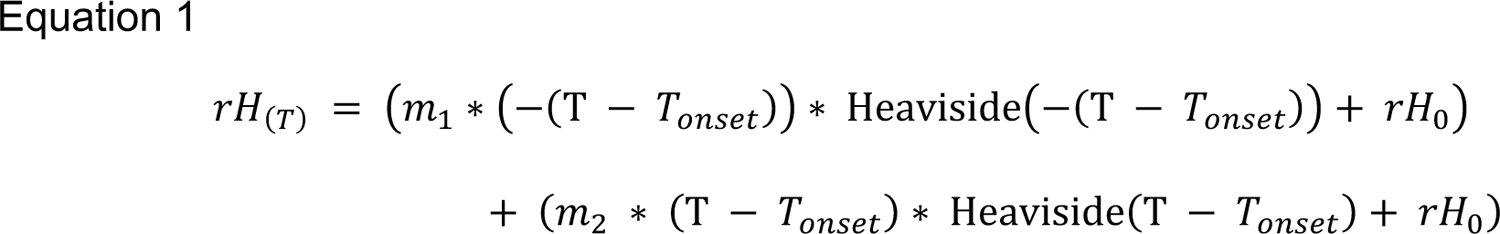

in which rH_(T)_ is the hydrodynamic radius at the measured temperature (T), rH_0_ is the extrapolated hydrodynamic radius at the beginning of the temperature range and m_1_ and m_2_ are the slopes of the linear functions before and after the assembly temperature (T_onset_), respectively. The two linear functions are concatenated using the Heaviside step function for conditional analysis of T < T_onset_ and T > T_onset_. Data analysis was with the R/Rstudio software package.

### Microscopy of heat-induced Ded1p assemblies

Fluorescence microscopy images of 4 μM GFP-labelled Ded1p and orthologs were taken with an Eclipse Ti2-E (Nikon, Japan) equipped with a Prime 95B 25 mm camera (Photometrics, USA) and using Plan Apo VC 60x/1.20 WI objective (Nikon, Minato, Japan) or with an Eclipse Ti (Nikon, Japan), equipped with CSU-X scan head (Yokogawa, Musashino, Japan) and an iXON 897 EMCCD camera (Andor, Belfast, Northern Ireland) using a 60x/1.2 Plan Apochromat water objective (Nikon, Japan). Proteins were heated to the indicated temperatures for 15 min in 20 mM PIPES/NaOH, 150 mM KCl and 0.5 mM TCEP in the presence or absence 100 ng/ml *in vitro* transcribed RNA. After heating, Ded1p assemblies were transferred to a medium binding Greiner µClear 384 well plate (Greiner Bio-One GmbH, Frickenhausen, Germany). The plates were briefly spun to allow sedimentation of the assemblies before imaging at room temperature.

### In vitro transcription and RNA purification

An 800-nt RNA construct containing the Ded1p-dependent 5’UTR of *SSK2*, the open reading frame of the luciferase gene and a poly(A)-tail was *in vitro* transcribed (Iserman et al., 2020). Template DNA for the *in vitro* transcription reaction was amplified by PCR using Phusion polymerase (Thermo Fisher Scientific, Waltham, USA) and RNA was synthetized using T7 RiboMAX kit (Promega, Madison, USA) during a 2 h incubation step according to the manufacturers’ protocol. The RNA was polyadenylated using the Poly(A) Tailing kit (Thermo Fisher Scientific, Waltham, USA) for 30 minutes and by following the manufacturer’s instructions. The RNA was precipitated with lithium chloride at −20°C, washed with 70% ethanol and resuspended in nuclease-free water. RNA concentrations were determined with Bioanalyzer 2100 (Agilent, Santa Clara, USA) with the RNA 6000 Nanokit (Agilent, Santa Clara, USA).

### Nano differential scanning fluorimetry (nanoDSF) and light scattering

NanoDSF and light scattering of purified proteins were measured with a Prometheus NT.48 (Nanotemper, Munich, Germany), equipped with back-reflection optics or with a Prometheus Panta (Nanotemper, Munich, Germany), equipped with back-reflection and DLS optics. Samples were assessed at 5-10 μM protein in 200 mM KCl, 32 mM PIPES pH 7.4 and 0.4 mM TCEP in high sensitivity capillaries, unless indicated differently. The samples were heated at a rate of 1°C/min from 20°C-95°C. For experiments in which DLS was acquired in addition to nanoDSF, the heating rate was 0.5°C/min and the cumulant radius was summarized in 1°C bins during analysis. For all experiments, the excitation was at 280 nm at 100% excitation power and emission intensities were recorded at 330 nm and 350 nm. When applicable, the excitation power of the DLS laser was 100%. The data were fitted using equation 2,

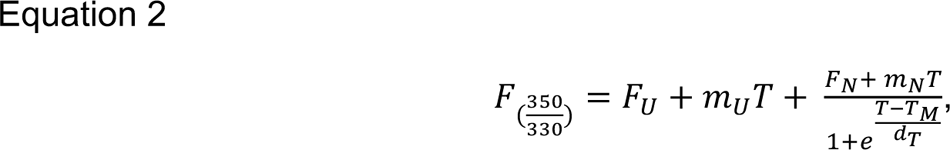

in which the fluorescence ratio (F_(350/330)_) was analyzed as a function of the temperature (T) with the parameters F_N_ and F_U_ (fluorescence ratio of the folded and unfolded states, respectively), m_N_ and m_U_ (drifts of the native and unfolded baselines, respectively) and T_M_ (transition midpoint) and d_T_ (slope of the transition midpoint) Light scattering data was analyzed using equation 3,

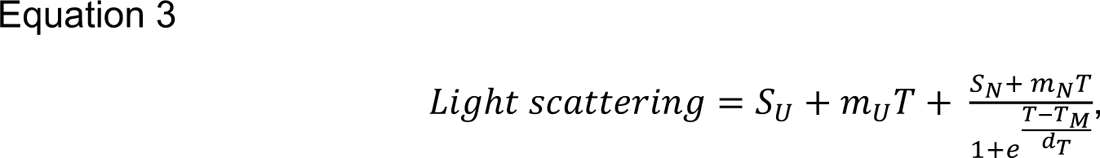

in which the light scattering signal was analyzed as a function of the temperature T with the parameters S_N_ and S_U_ (light scattering signal of the unassembled and assembled protein, respectively), the slopes of the baselines mN and mU and apparent transition midpoint (TM) and slope of the transition midpoint (dT).

For visual purposes, the data was normalized using equation 4

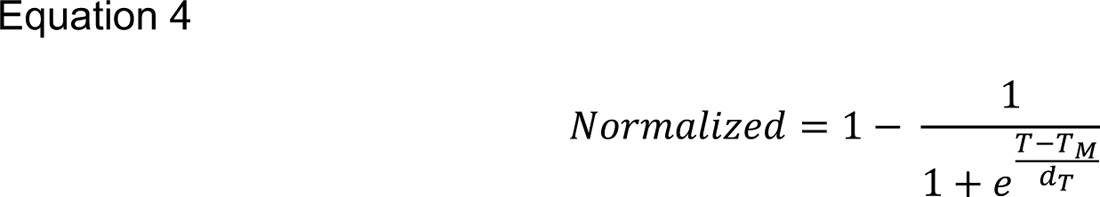

with the result parameter values for T_M_ and d_T_ derived from non-linear regression of the raw data using equation X for fluorescence ratio data and Y for light scattering data. T_onset_ was calculated using the equation 5

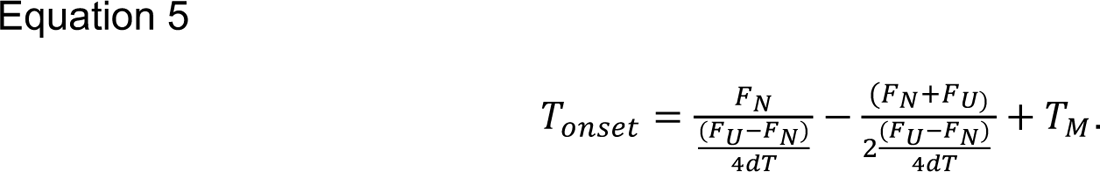

with the result parameters values F_N_, F_U_, T_M_ and d_T_ derived from non-linear regression of the raw data using equation X. For light scattering data, the result parameter values S_N_, S_U_, T_M_ and d_T_ were used. Data analysis and plotting was with the R/Rstudio software package.

### NMR Spectroscopy

The N-terminal region (residues 1-81, internal reference #2269), the first RecA domain (residues 82-369, internal reference 1739), the second RecA domain (residues 369-535, internal reference #1735), the helicase core region (residues 82-535, internal reference #1740) and the C-terminal IDR (residues 533-604, internal reference #2000) of Ded1p were cloned into a modified pET-vector that carries an N-terminal TEV protease cleavable His_6_-tag. Proteins were expressed overnight at 20°C in *E. coli* using minimal medium that was supplemented with ^15^N ammonium chloride and/ or ^13^C_6_ glucose (for NMR visible proteins) or using LB medium (for NMR invisible proteins). The cells were collected by centrifugation (20°C, 20 minutes, 6.000 g), lysed by sonication in buffer NMR1 (25 mM NaPO_4_ pH 7.4, 1 M NaCl, 1 mM DTT, 10 mM imidazole) that was supplemented with 0.1% Triton, 1 U/mL DNAse1 and 0.1 mg/mL lysozyme, after which the cell debris was removed by a second centrifugation step (4°C, 30 minutes, 35.000 g). The supernatant was loaded onto a gravity flow Ni-NTA column, the column was washed with 5 column volumes buffer NMR1, 10 column volumes of buffer NMR2 (25 mM NaPO_4_ pH 7.4, 2 M NaCl, 1 mM DTT) and the bound protein was eluted with buffer NMR3 (25 mM NaPO_4_ pH 7.4, 1 M NaCl, 1 mM DTT, 300 mM imidazole). Overnight, the eluted proteins were dialyzed into buffer NMR4 (20 mM HEPES pH 7.3, 500 mM NaCl, 1 mM DTT) in the presence of 1 mg TEV protease. The cleaved affinity tag was removed by a second gravity flow Ni-NTA column using buffer NMR4, after which the protein was applied to a size exclusion column (HiLoad 16/600 Superdex 75) that was equilibrated in buffer NMR5 (20 mM HEPES pH 7.3, 125 mM NaCl, 1 mM DTT). The fractions that contained the pure protein were collected, concentrated, and supplemented with 5% D_2_O. The NMR samples contained between 50 µM and 1.5 mM protein.

NMR spectra were recorded on Bruker Avance NEO NMR spectrometers that operate at 500 or 800 MHz proton resonance frequency and that are equipped with nitrogen (500) or helium (800) cooled TCI probeheads. The backbone resonances of the Ded1p N-terminal IDR and the Ded1p RecA2 domain were assigned using standard 3-dimensional TROSY (Transverse relaxation optimized spectroscopy) based (Pervushin et al., 1997) HNCO, HN(CA)CO, HNCACB, HN(CO)CACB and (H)CC(CO)NH spectra that were processed using the NMRPipe/NMRView software package (Delaglio et al., 1995) and analyzed using CARA (R. Keller, The computer aided resonance assignment tutorial, 2004).

## Key Resources Table

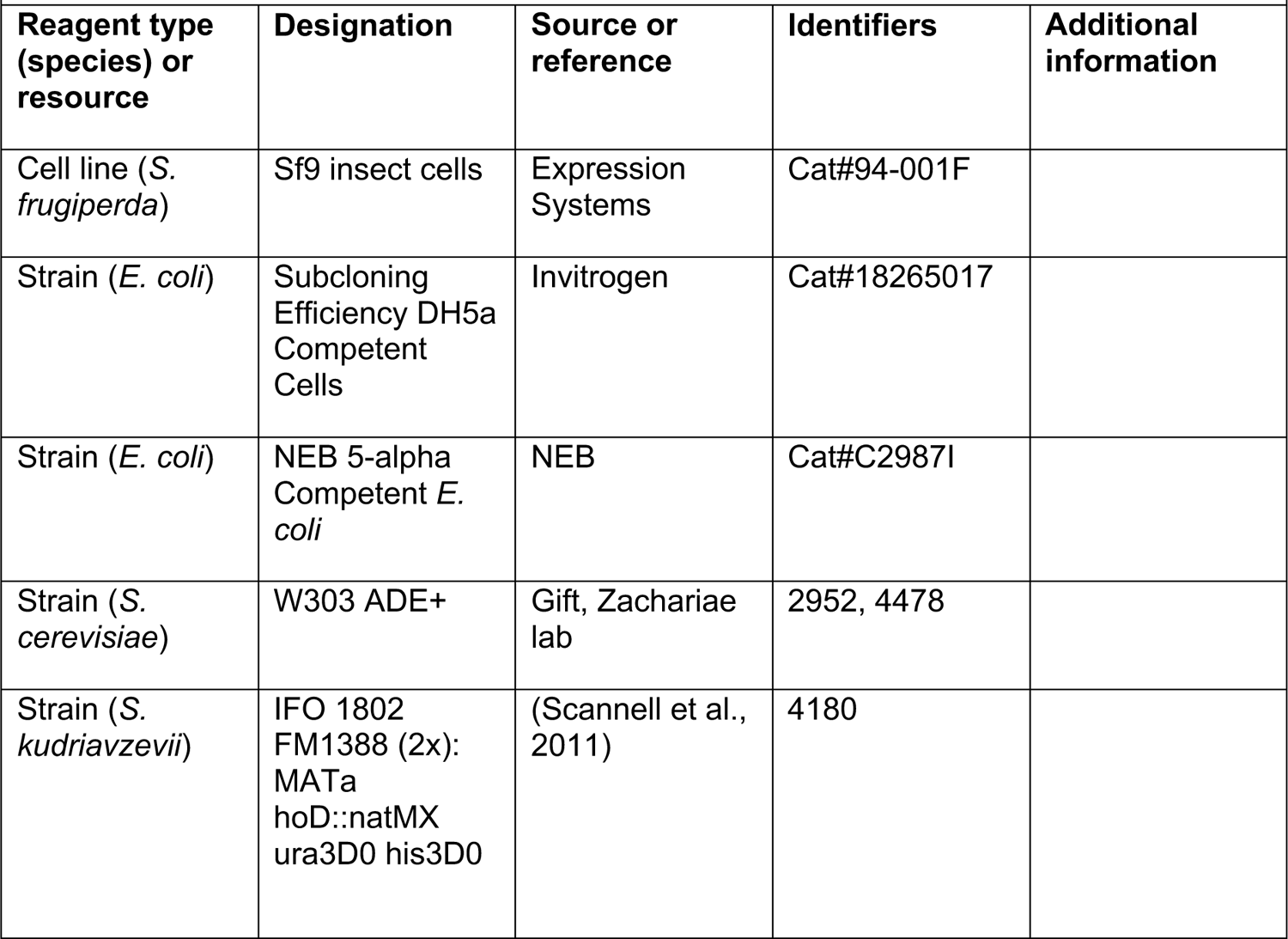
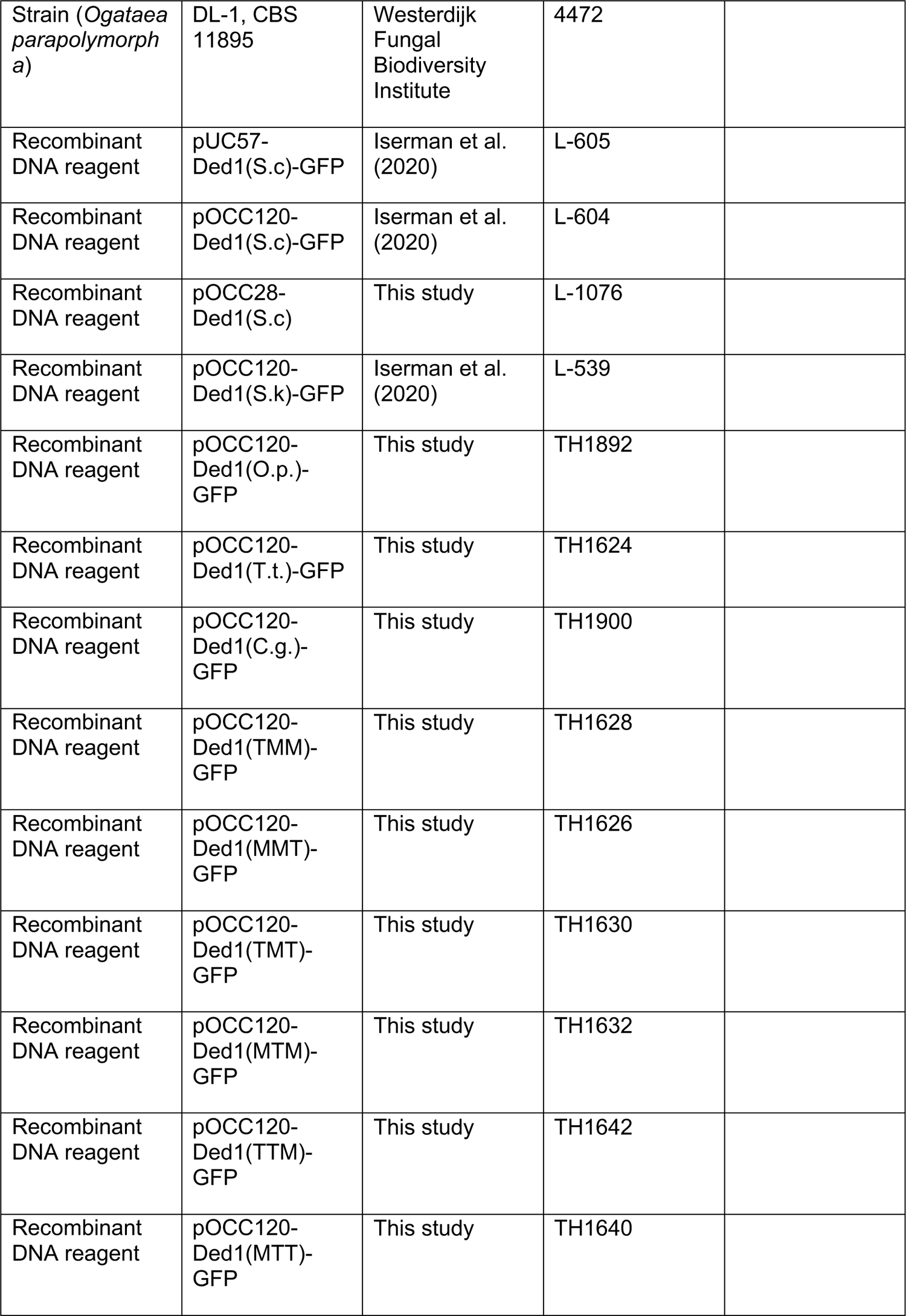
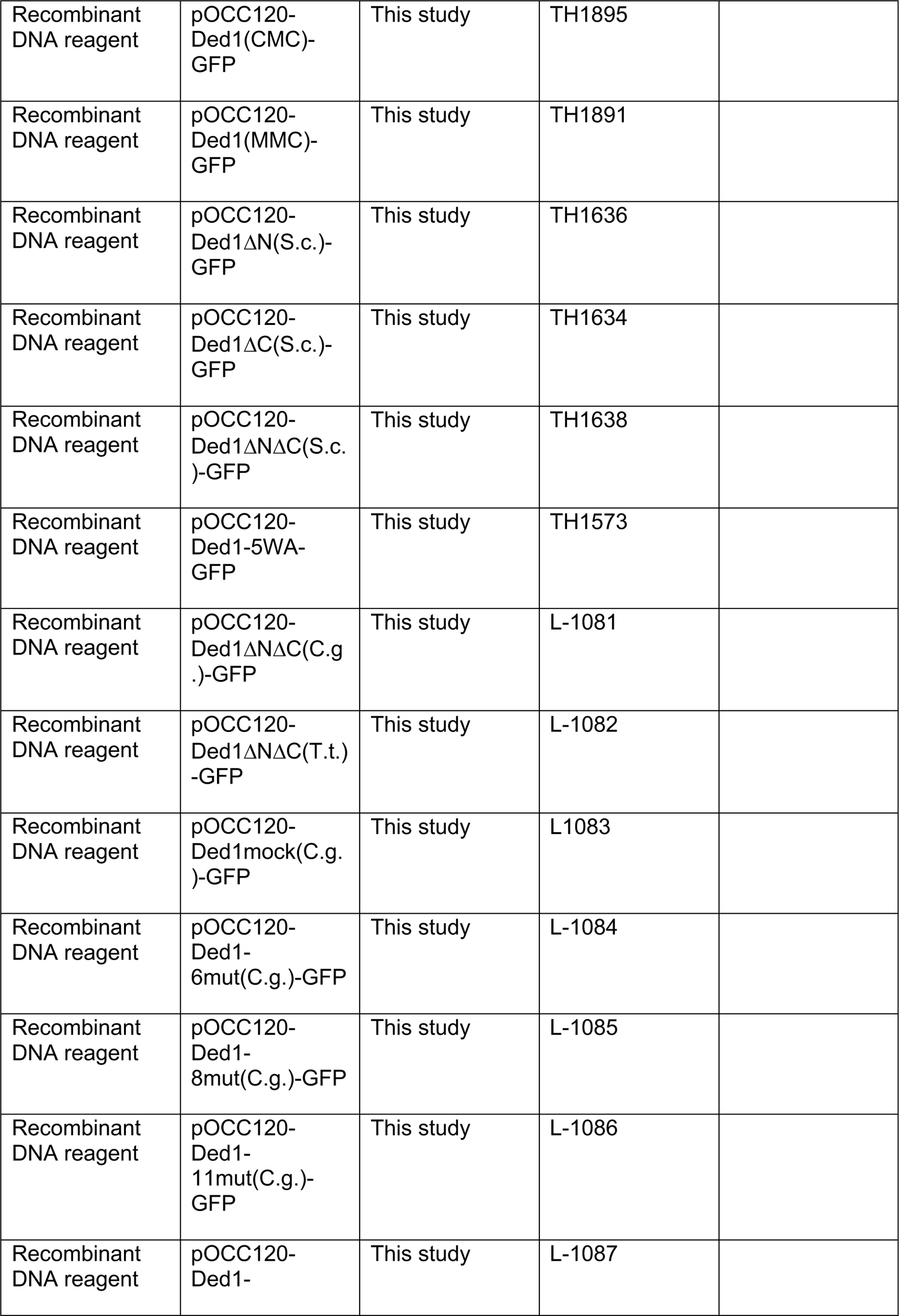
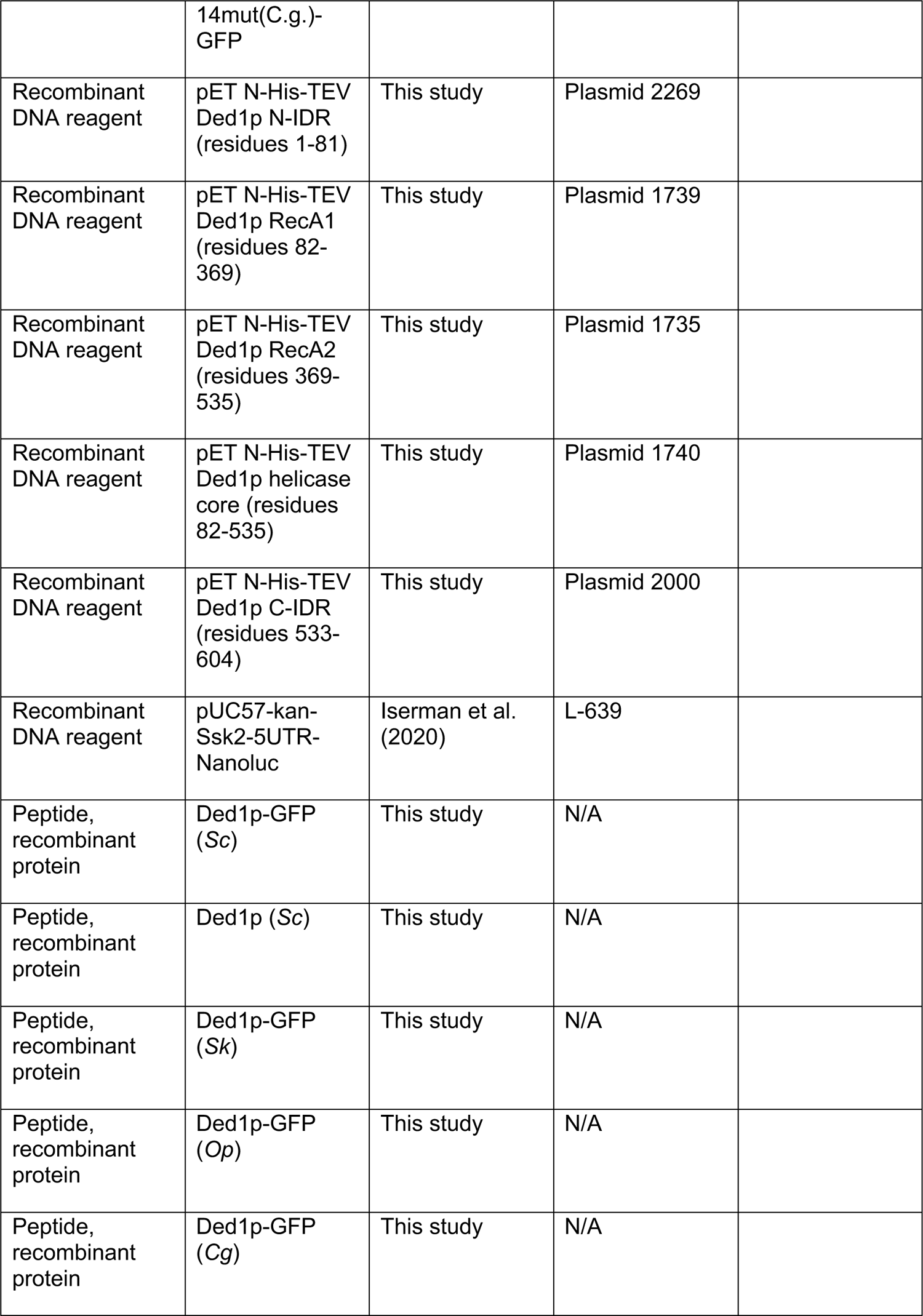
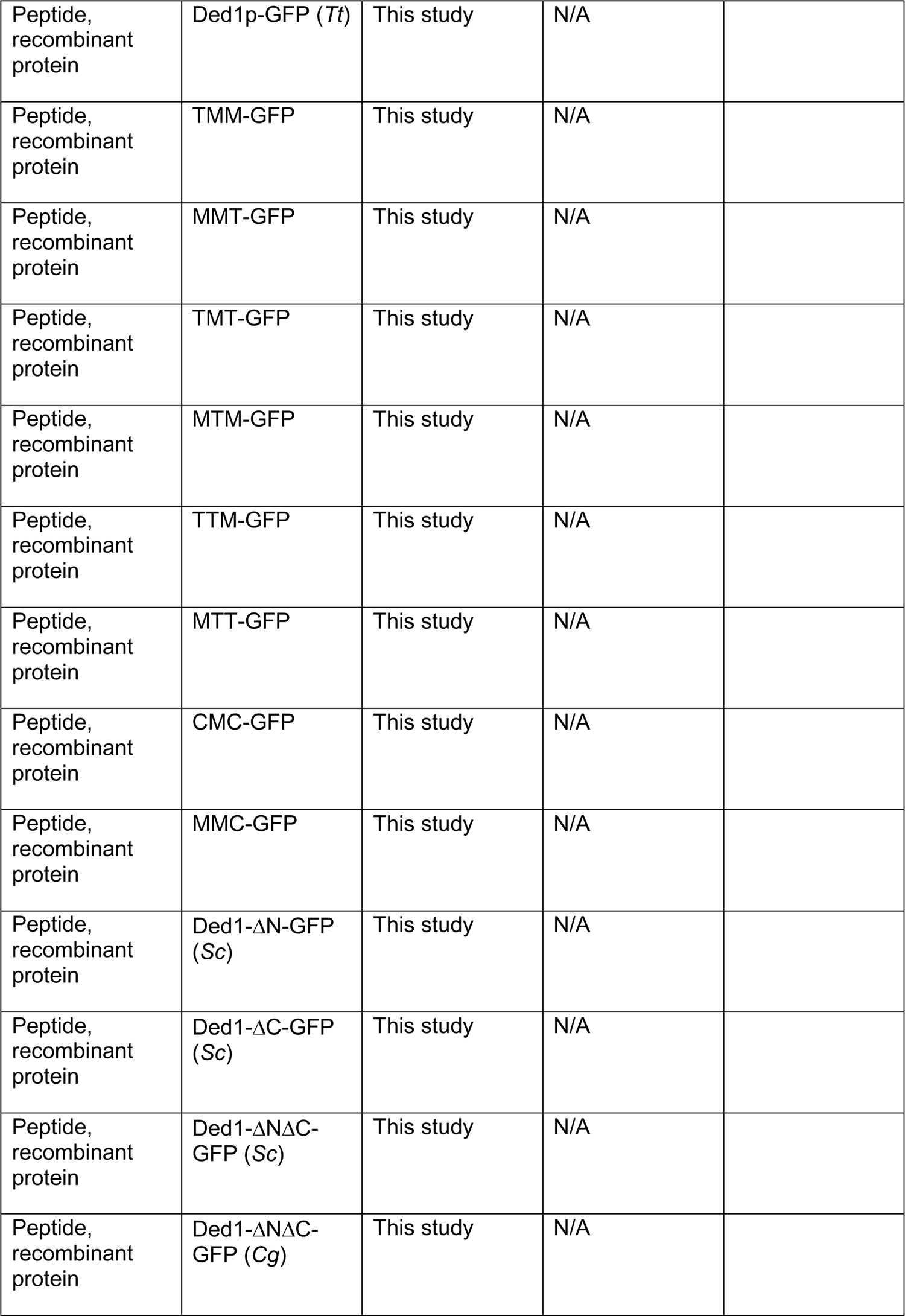
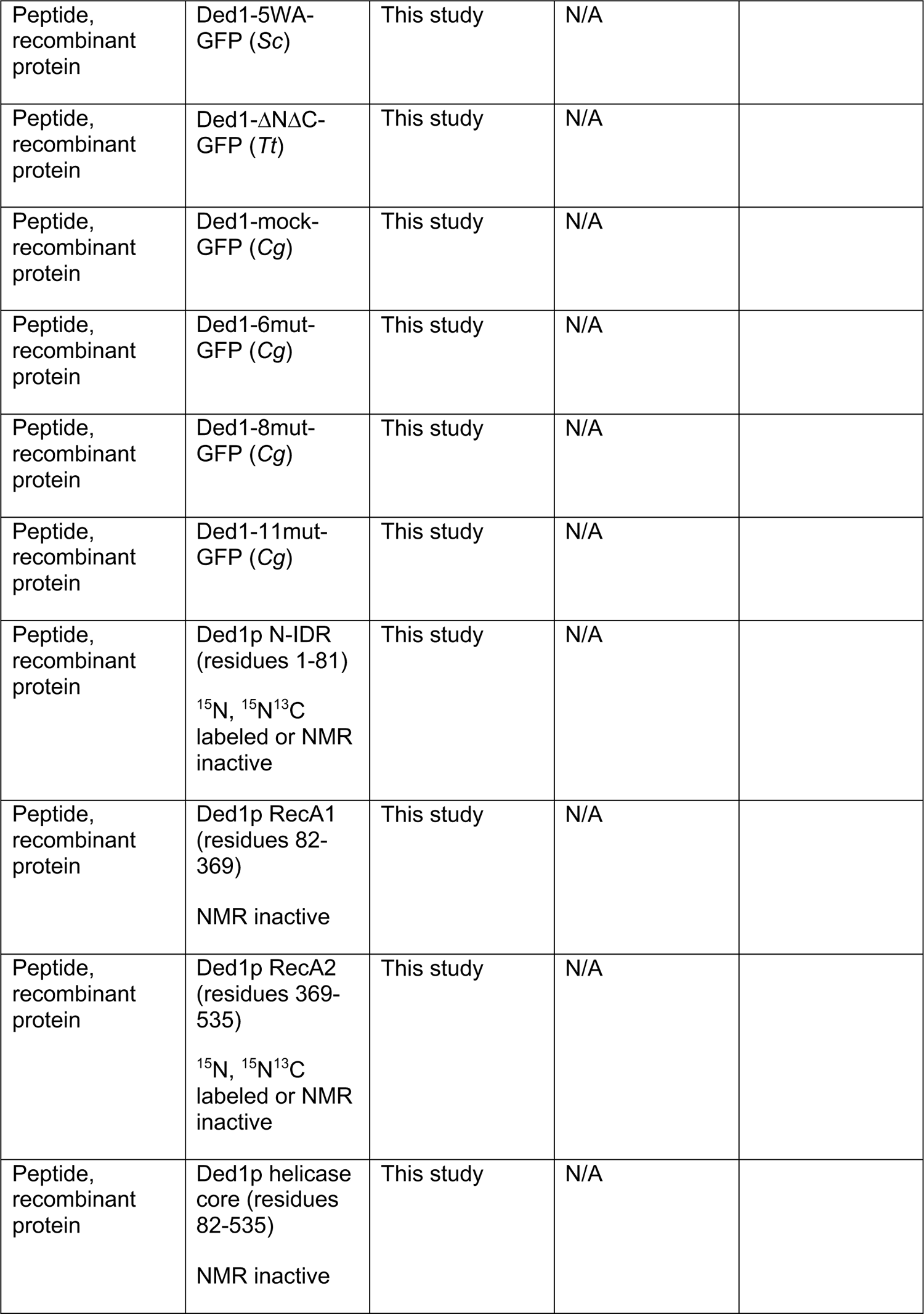
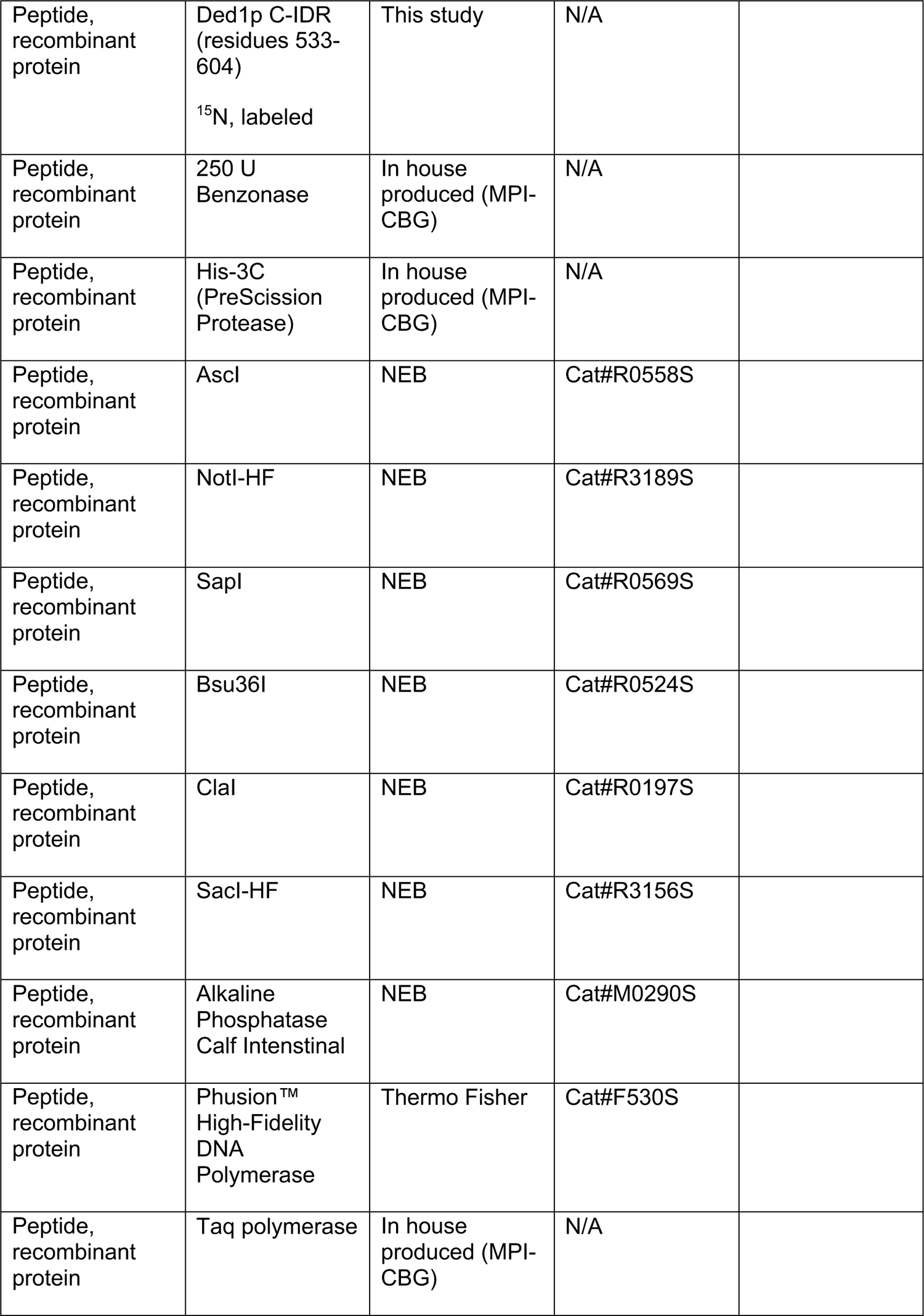
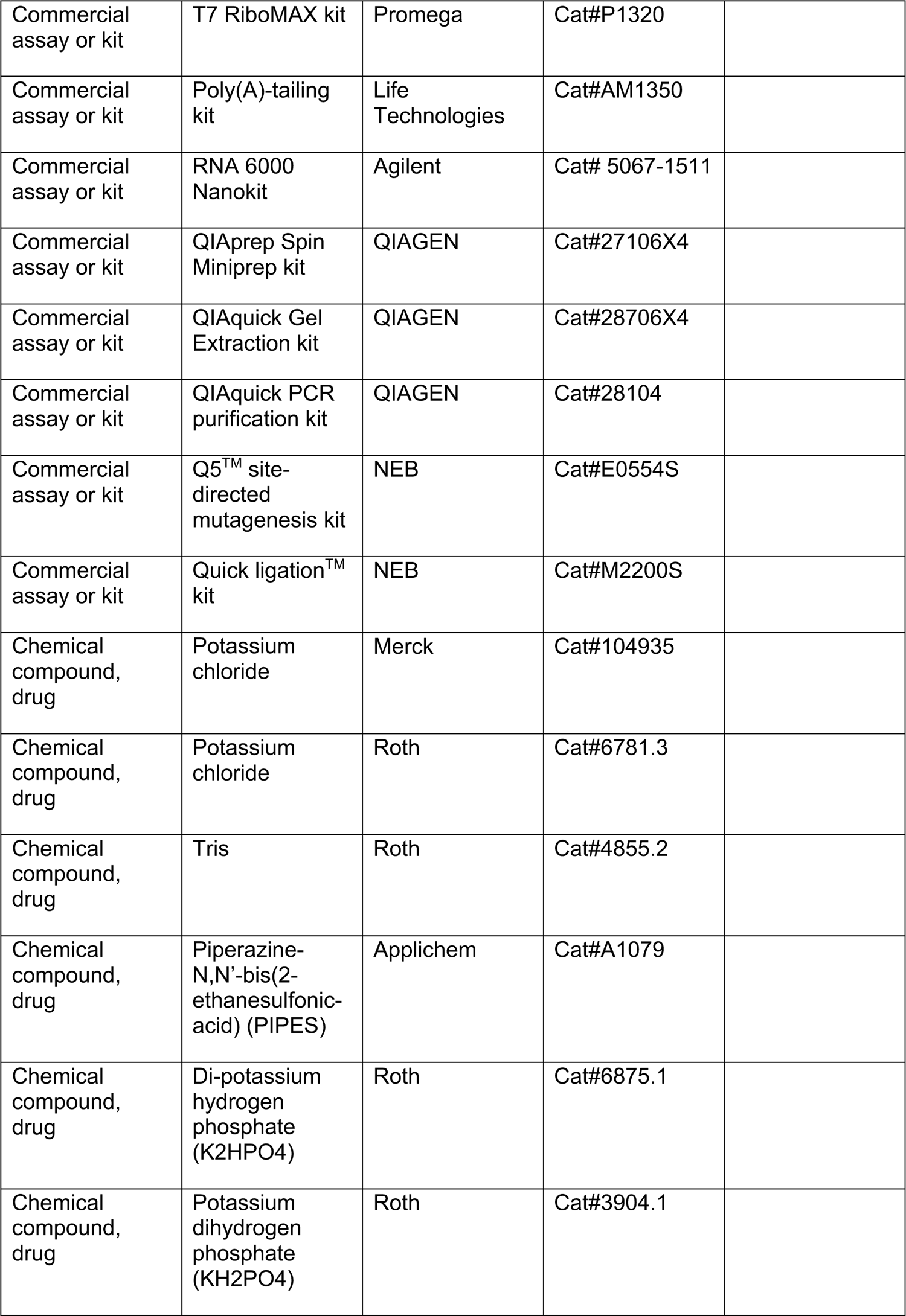
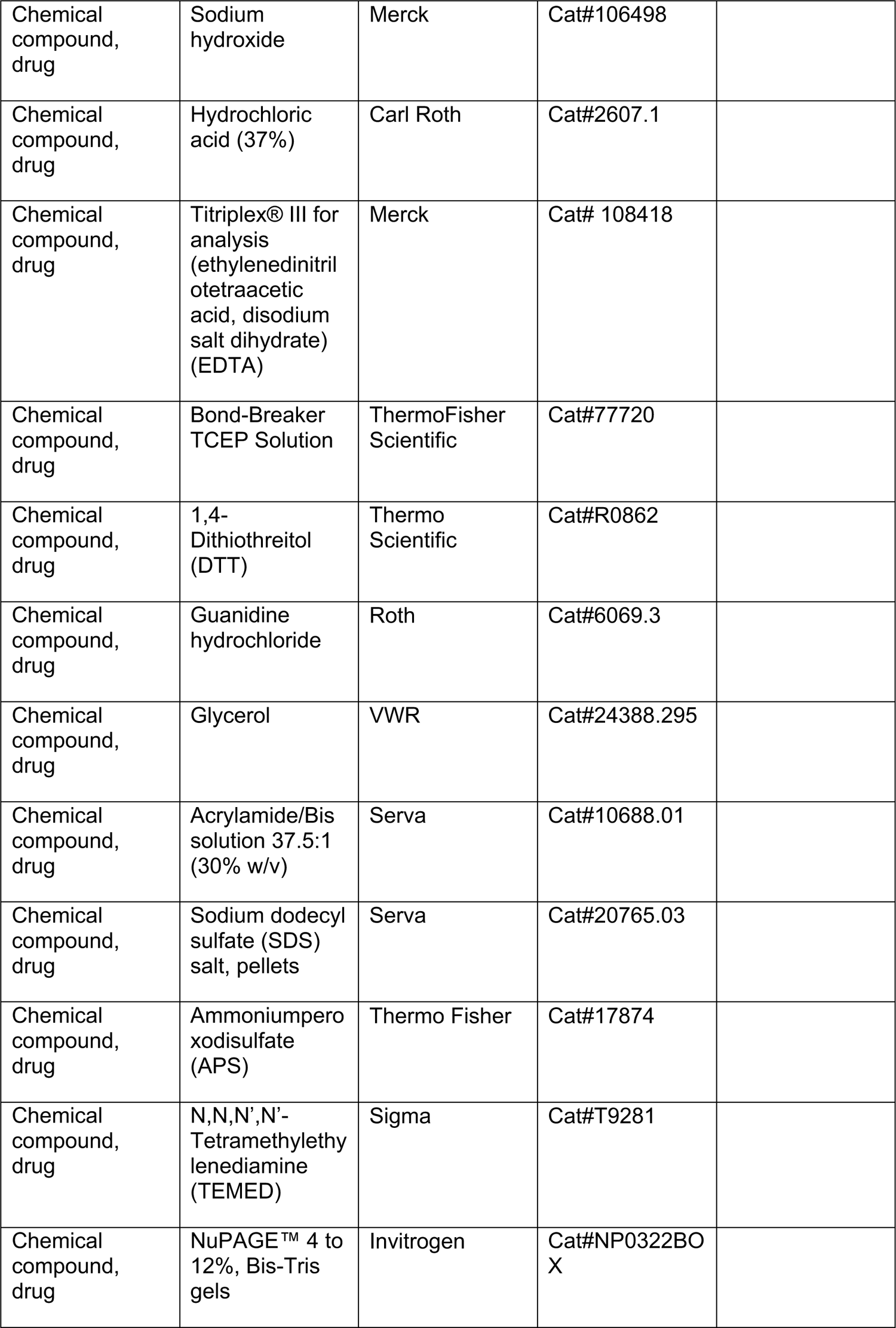
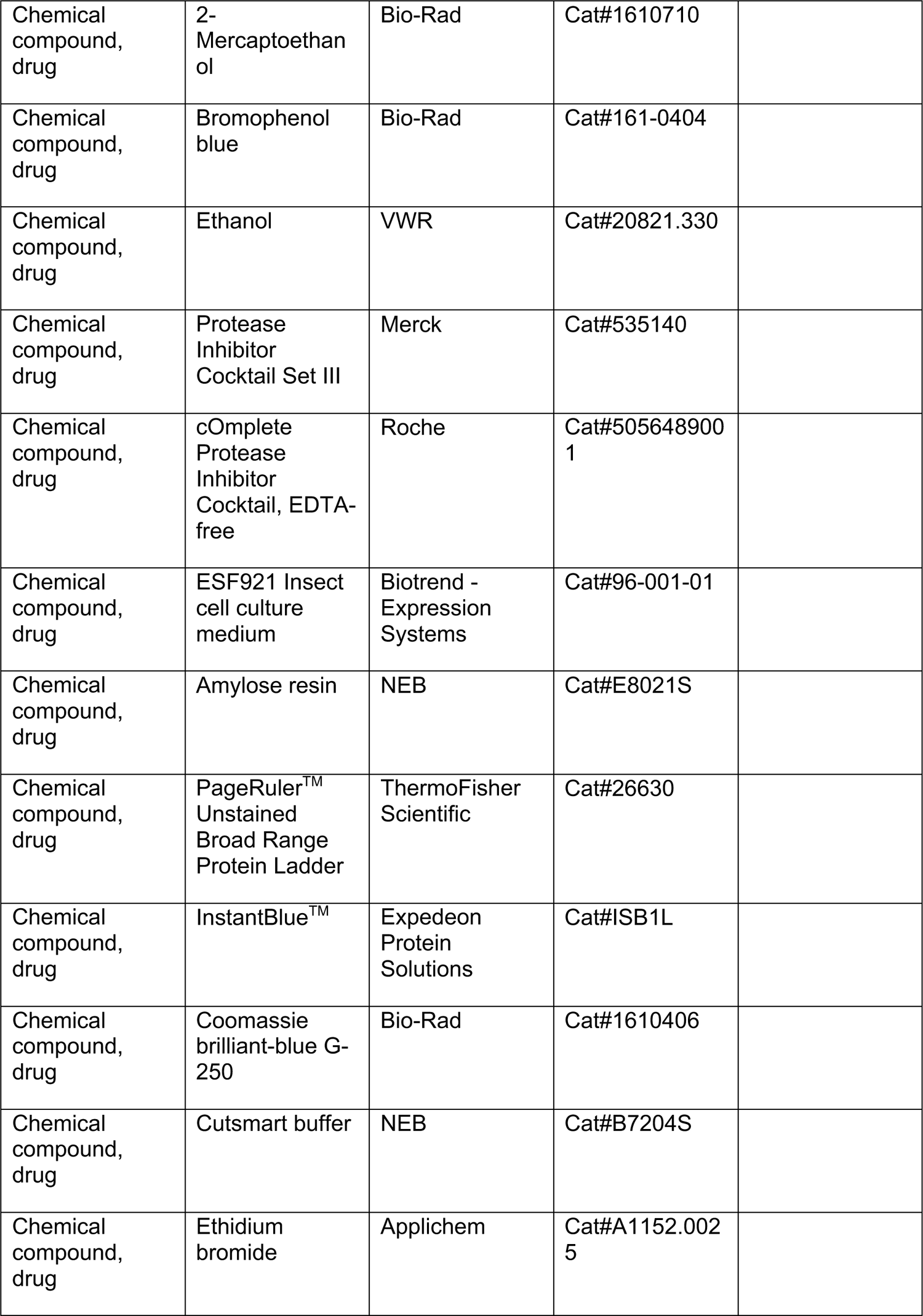
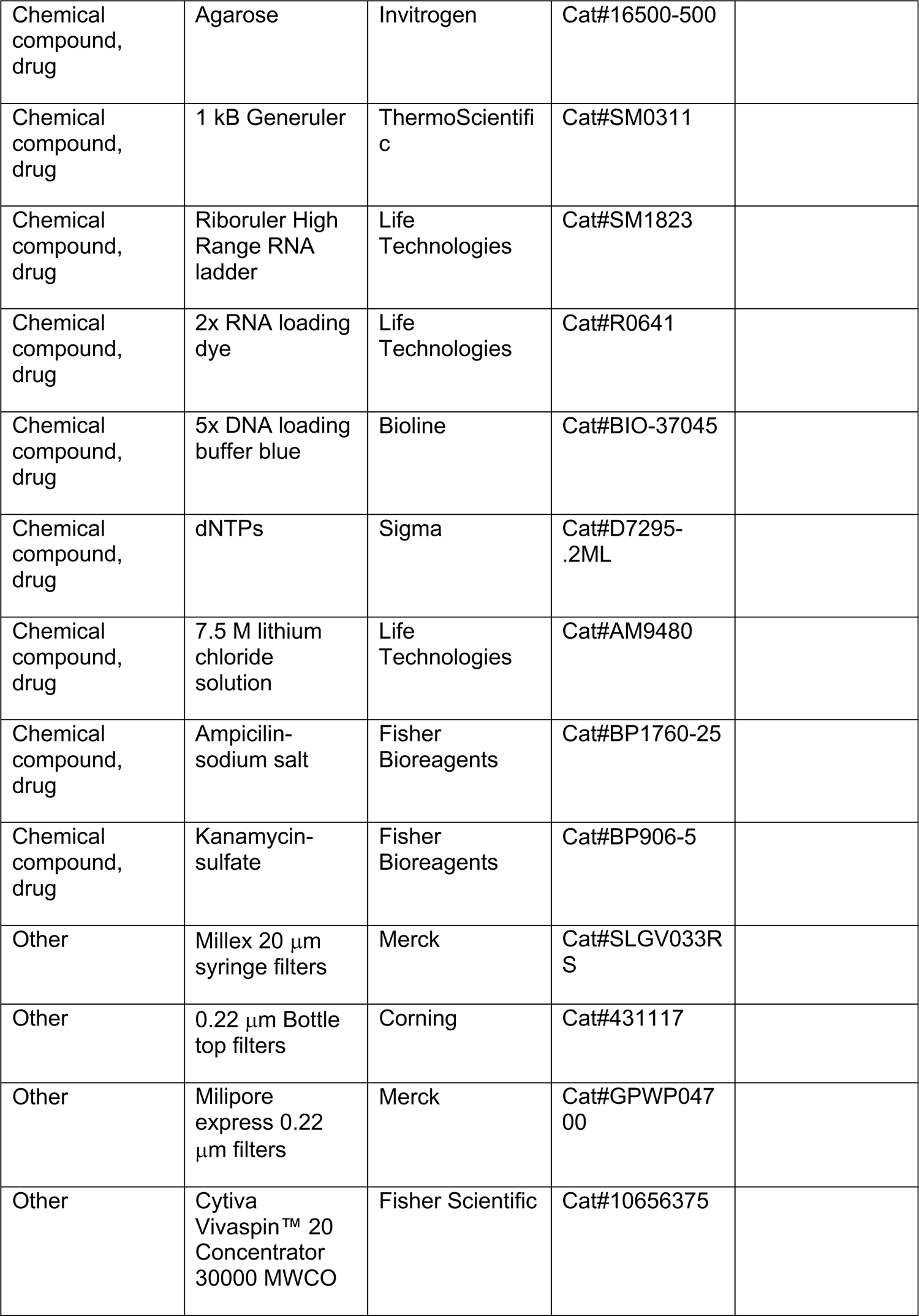
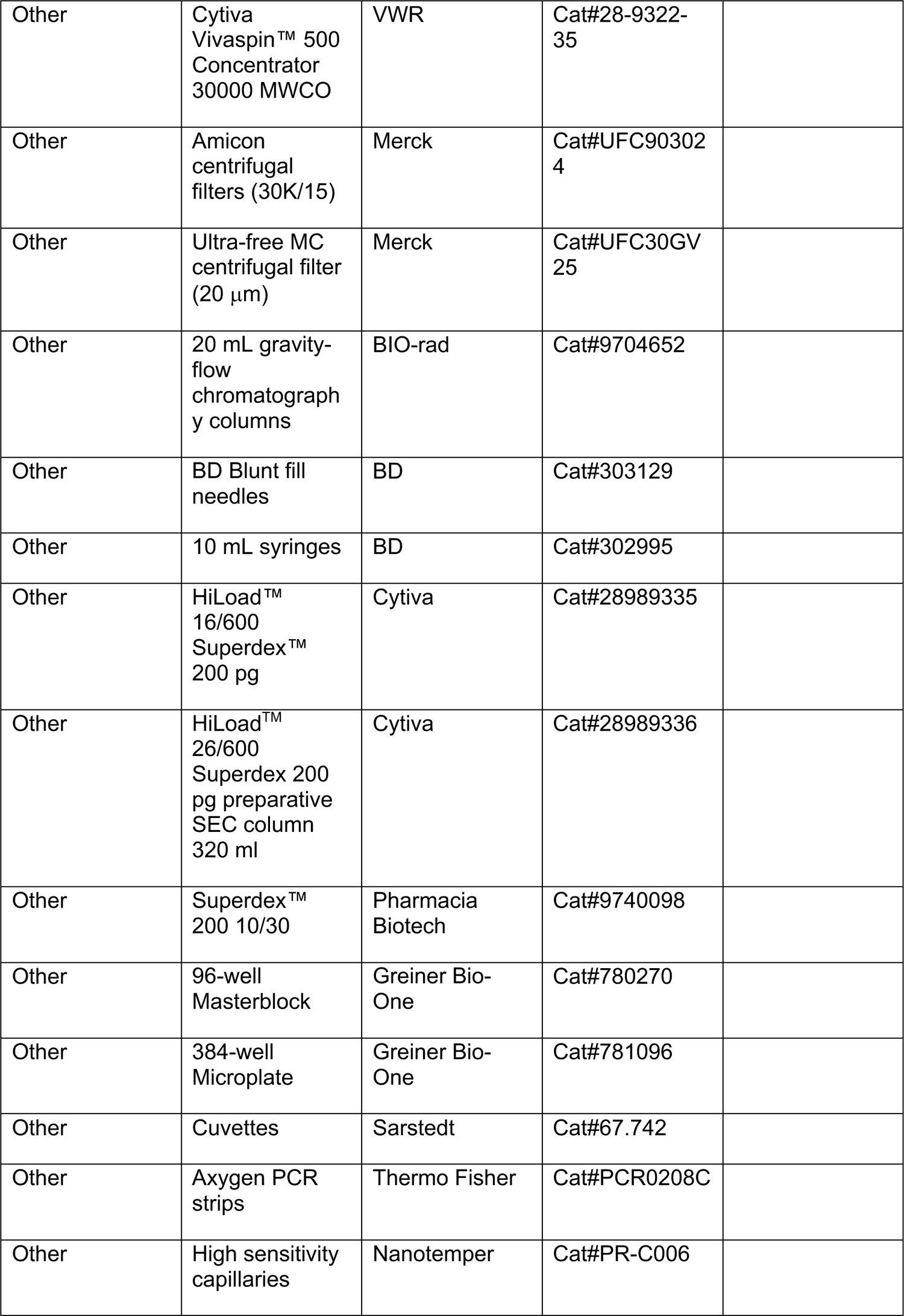
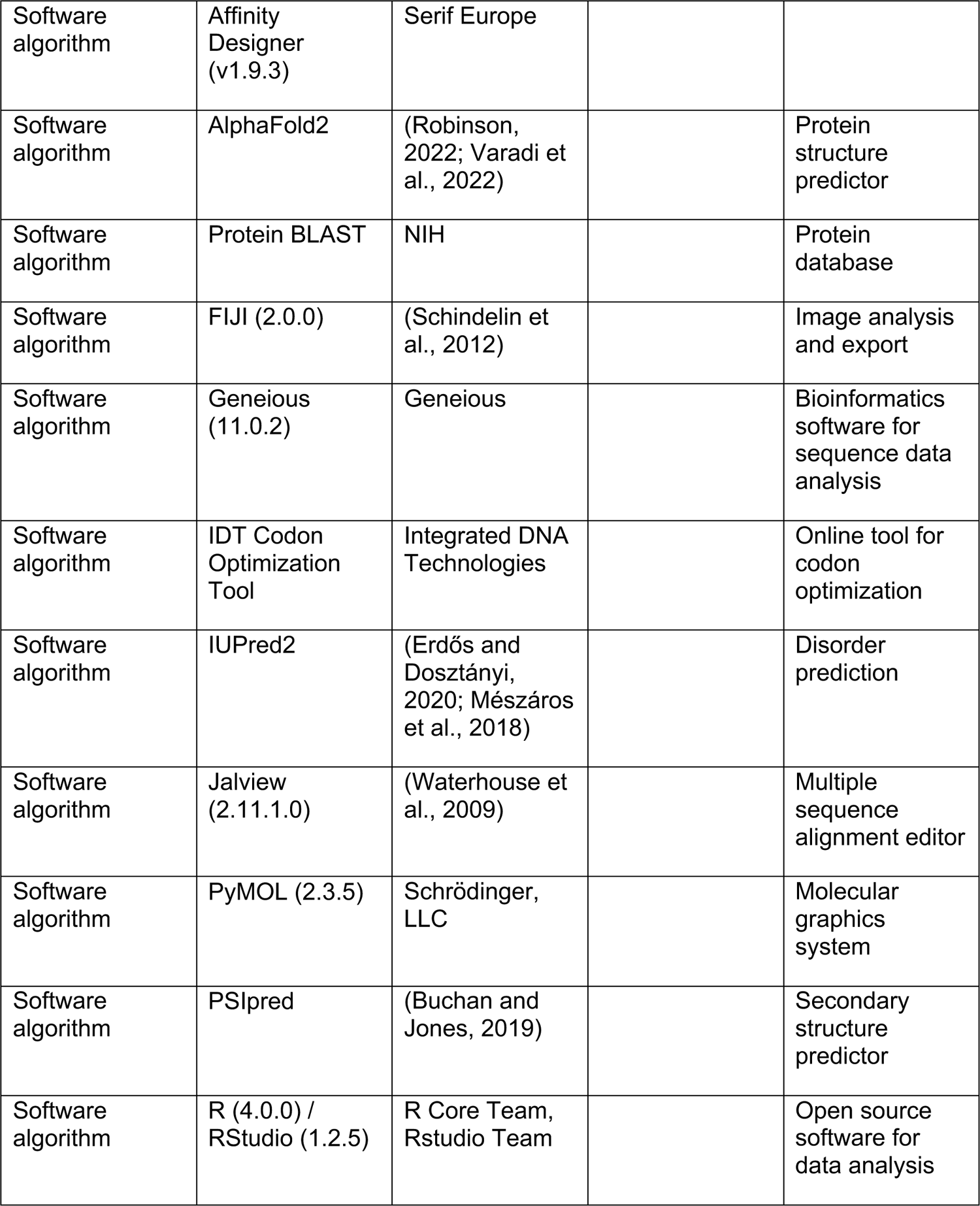

## Key Resources Table - Sequence based reagents

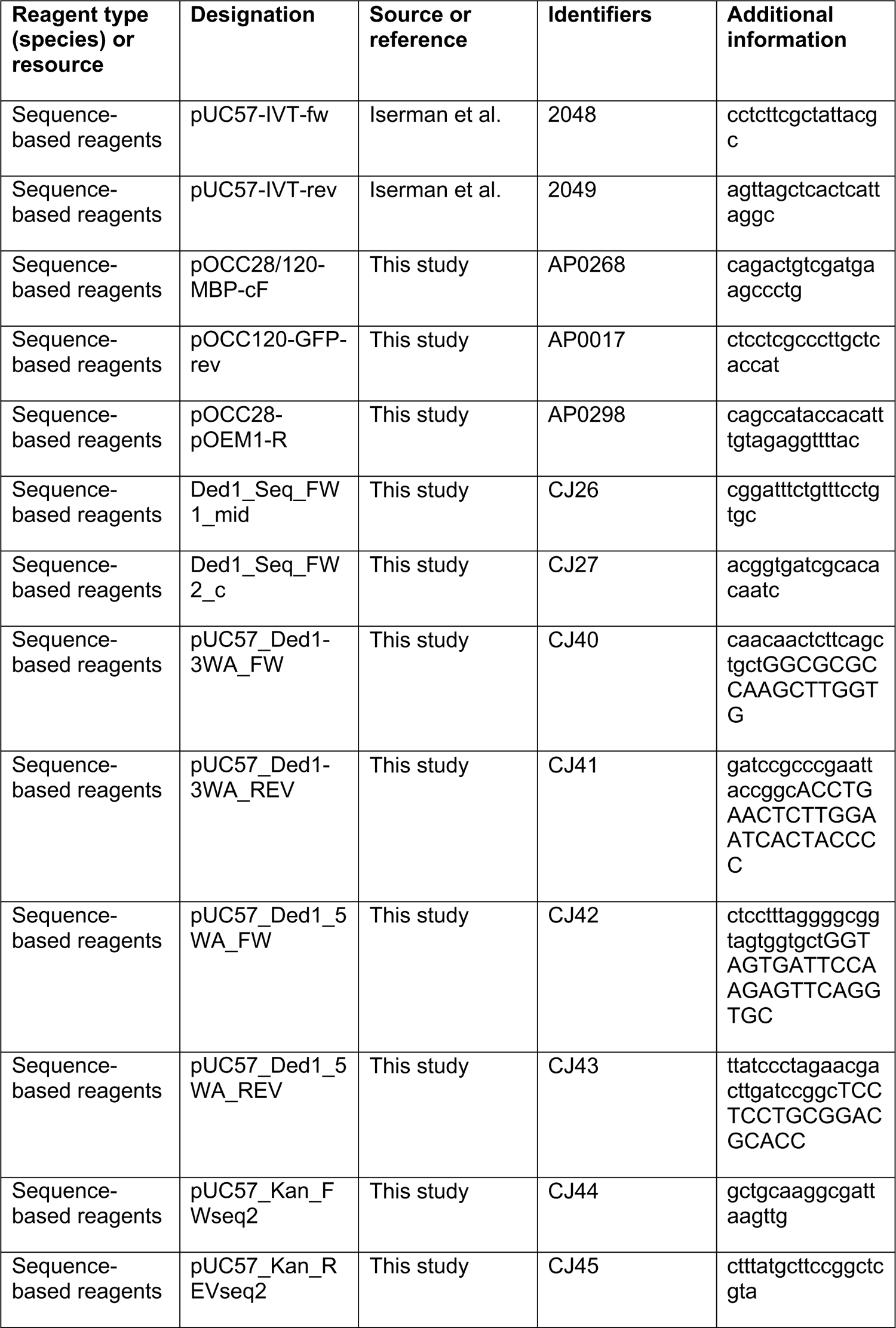
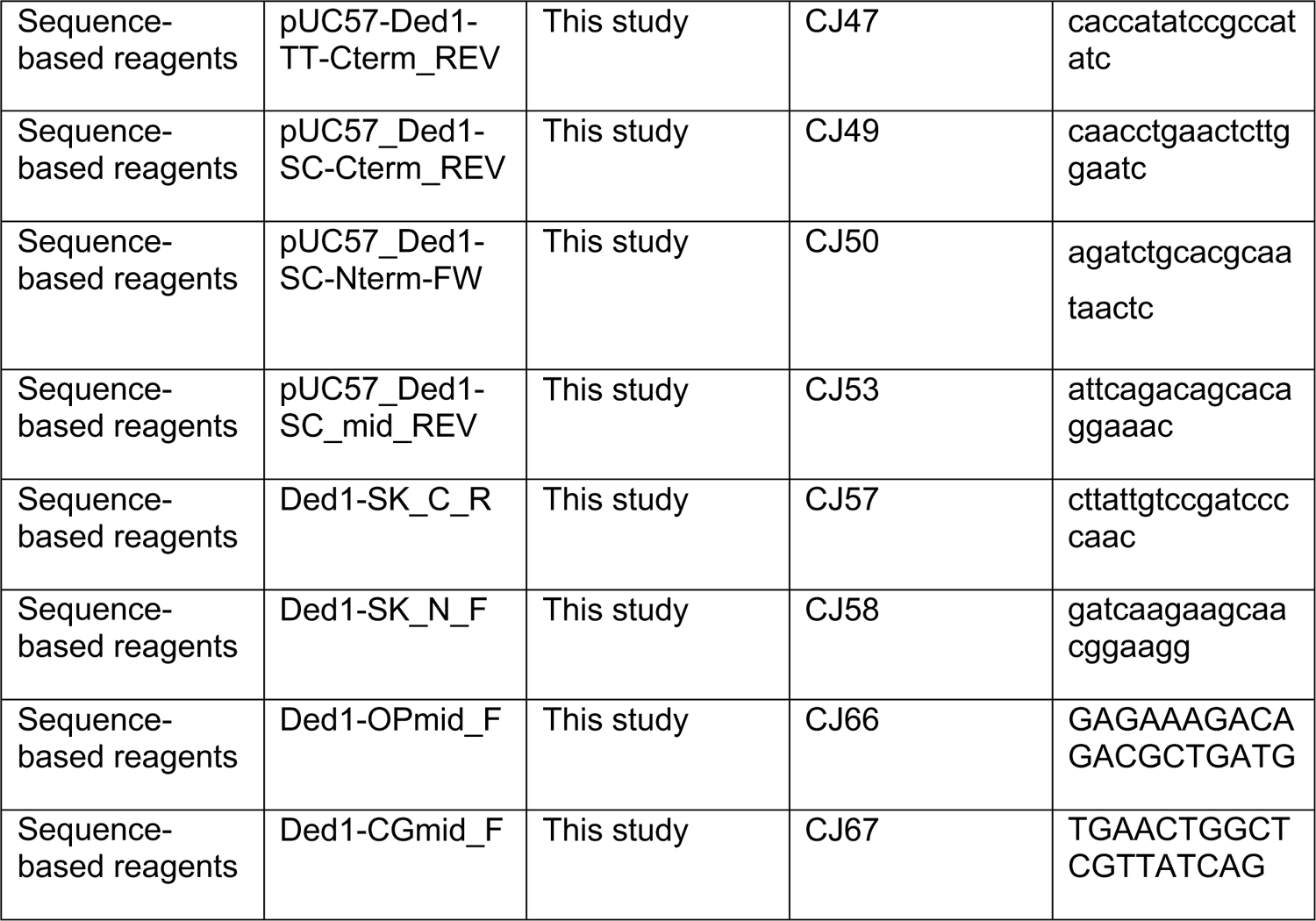

---

**Supplemental figure 1.**
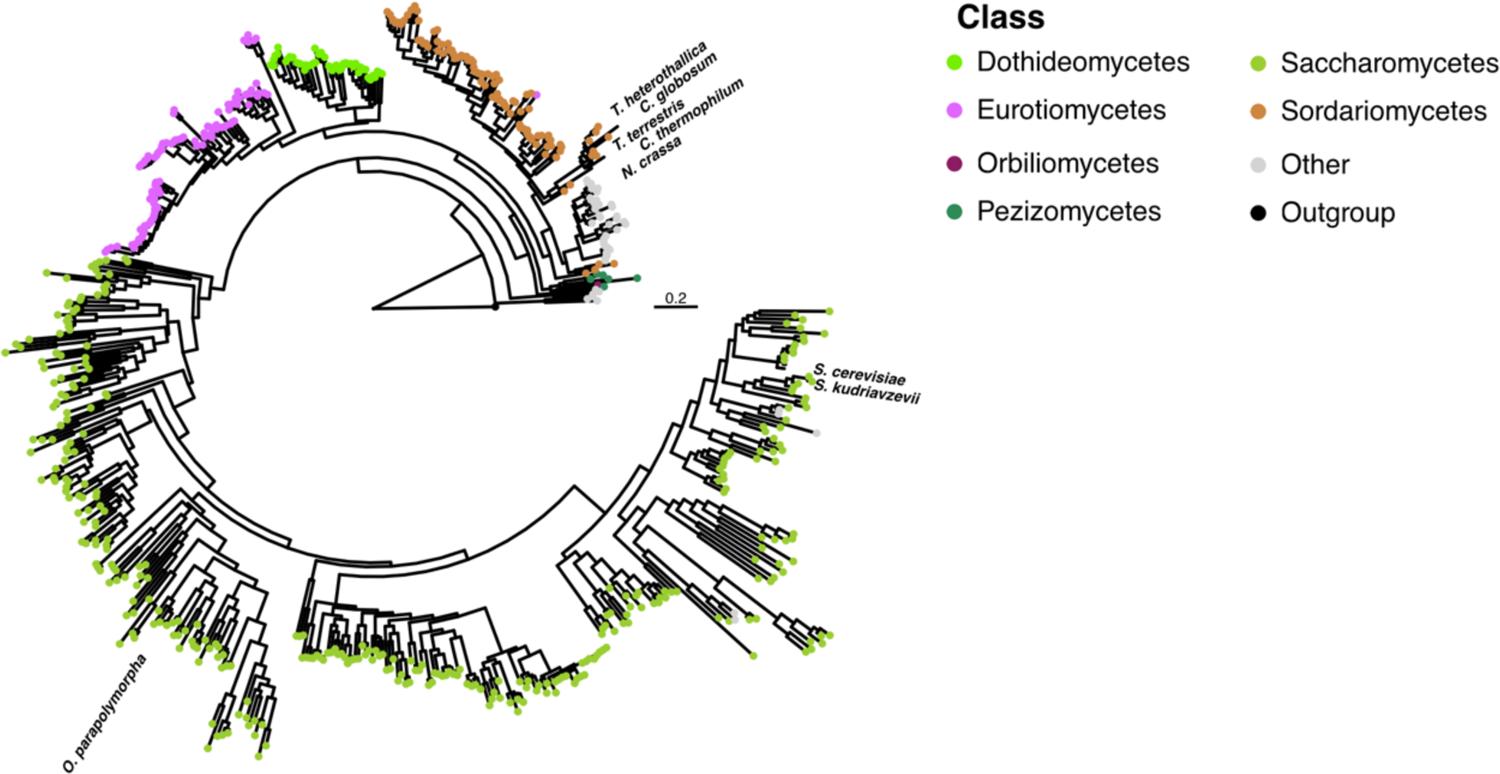
Phylogenetic tree based on fungal Ded1p orthologs. Scale = 0.2 substitutions per site.

**Supplemental figure 2.**
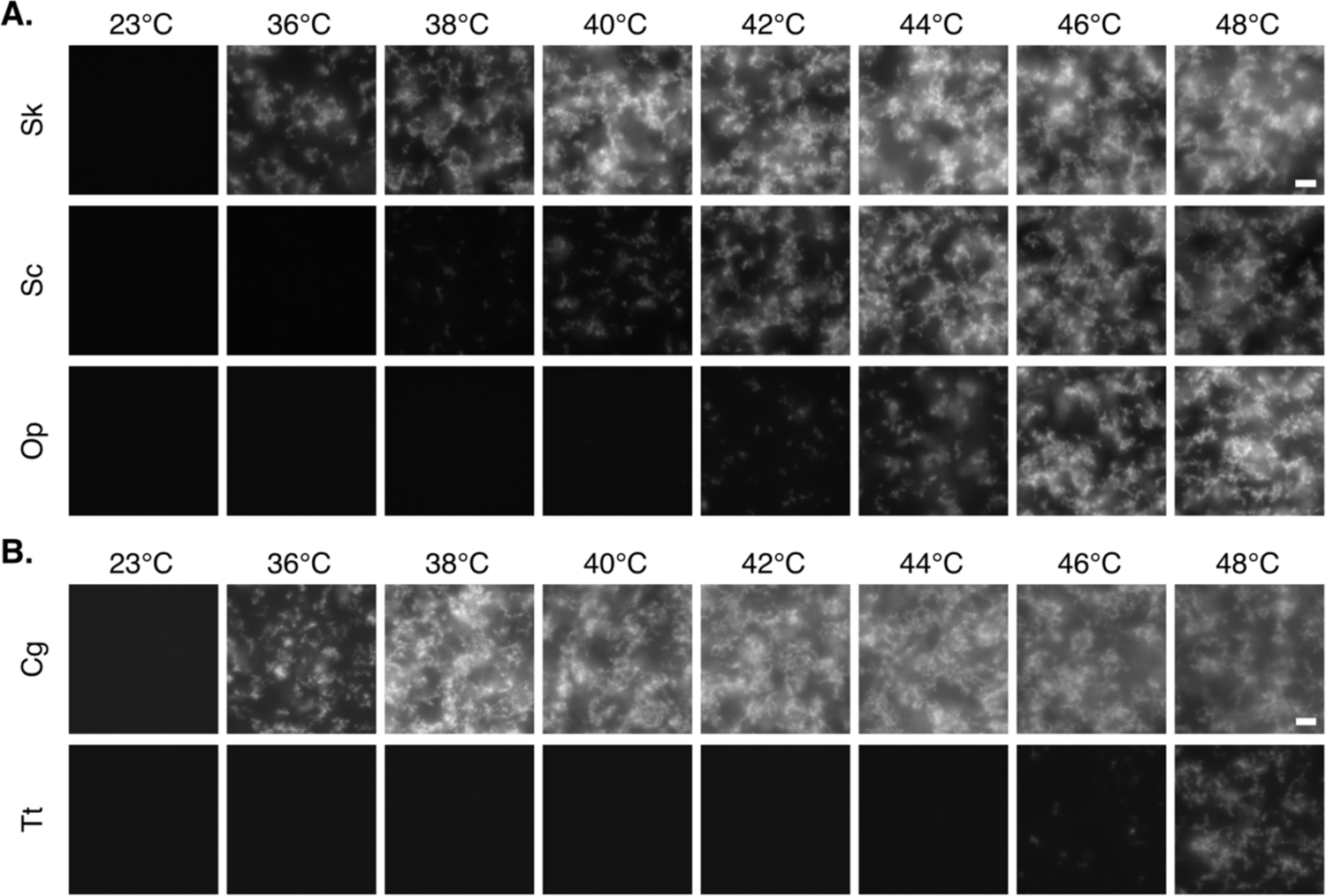
A. Representative fluorescence images of GFP-labelled Ded1p orthologs from *S. cerevisiae* (Sc)*, S. kudriavzevii* (Sk) and *O. parapolymorpha* (Op) heated to different temperatures. Scale bar = 10 μm. **B.** Representative fluorescence images of GFP-labelled Ded1p orthologs from *C. globosum* (Cg) and *T. terrestris* (Tt) heated to different temperatures. Scale bar = 10 μm.

**Supplemental Figure 3.**
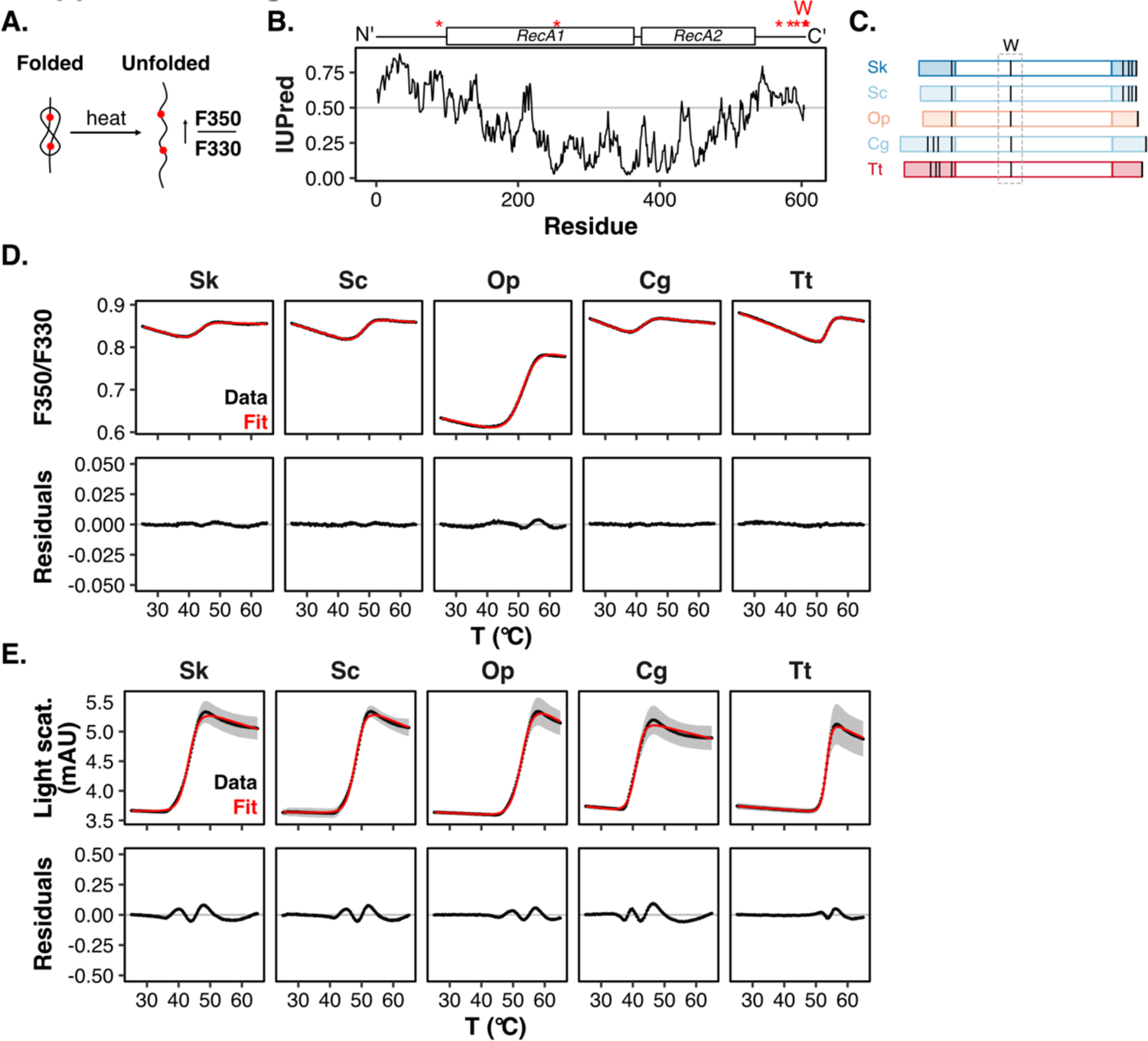
A. Schematic representation of thermal protein unfolding. Initially solvent-excluded tryptophan residues (red spheres) can become solvent-exposed upon unfolding, shifting the emission peak of Trp residues from λ = 330 to λ = 350. **B.** A domain structure of Ded1p from *S. cerevisiae*, highlighting the position of tryptophan residues (W), and a disorder prediction using IUPred (Dosztányi et al., 2005).**C.** Domain architecture of Ded1p orthologs from *S. cerevisiae* (Sc)*, S. kudriavzevii* (Sk)*, O. parapolymorpha* (Op), *C. globosum* (Cg) and *T. terrestris* (Tt), highlighting the distribution of tryptophan (W) residues with black bars. **D.** Change in F350/F330 as a function of temperature for 5 μM of the GFP-labelled Ded1p orthologs from *S. cerevisiae* (Sc)*, S. kudriavzevii* (Sk)*, O. parapolymorpha* (Op), *C. globosum* (Cg) and *T. terrestris* (Tt), showing the raw data (top panel, black), fitted data (top panel, red) and the residuals (bottom panel). Mean (solid lines), sd (grey ribbon), n = 6. **E.** Change in back-reflection light scattering as a function of temperature for 5 μM of the GFP-labelled Ded1p orthologs from *S. cerevisiae* (Sc)*, S. kudriavzevii* (Sk)*, O. parapolymorpha* (Op), *C. globosum* (Cg) and *T. terrestris* (Tt), showing the raw data (top panel, black), fitted data (top panel, red) and the residuals (bottom panel). Mean (solid lines), sd (grey ribbon), n = 6.

**Supplemental figure 4.**
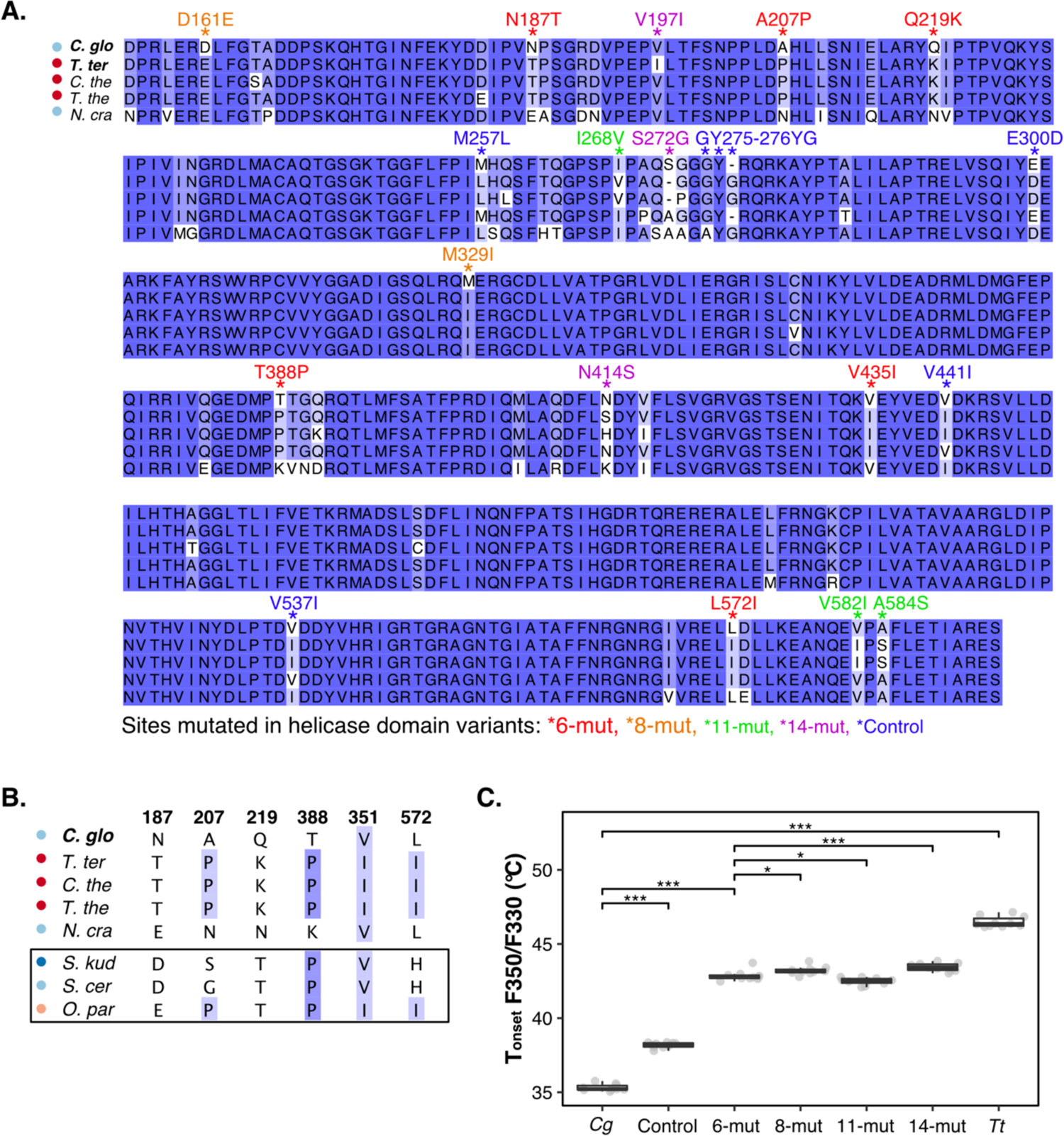
A. Alignment of the amino acid sequences for the helicase domains of mesophilic (*C. globosum, N. crassa*) and thermophilic (*T. terrestris, C. thermophilum* and *T. thermophilus*) Ded1p orthologs using MUSCLE. Residues mutated in the 6-mut, 8-mut, 11-mut and control variants are highlighted. **B.** Alignment of the six sites mutated in the variant “6-mut” across mesophilic and thermophilic fungi (see A), as well as in the budding yeast *S. cerevisiae, S. kudriavzevii* and *O. parapolymorpha*. **C.** Apparent T_onset_ of F350/F330 for 8 μM of the GFP-labelled helicase domains of *C. globosum* (C.g.) and *T. terrestris* (Tt) and variants (6-mut, 8-mut, 11-mut, 14-mut and Control) using nanoDSF (n = 10, *** p < 0.001 calculated using T-test).

**Supplemental Figure 5.**
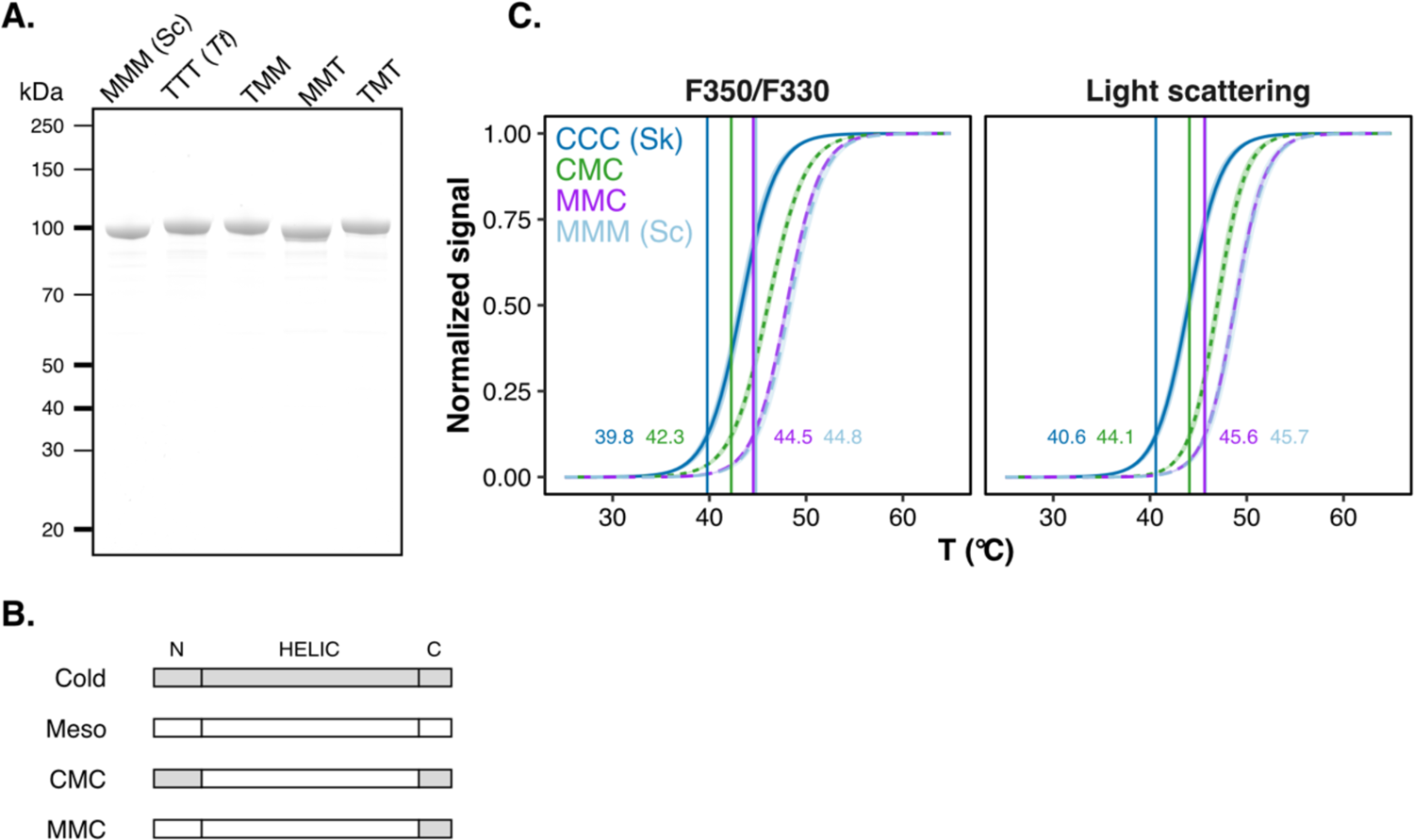
A. Coomassie stained SDS gel of GFP-labelled full length *S. cerevisiae* Ded1p, *T. terrestris* Ded1p and corresponding chimeric proteins (see Figure 2A). **B.** Domain architecture of Ded1p from *S. cerevisiae* (mesophile, meso) and *S. kudriavzevii* (cold-adapted, cold) and chimeric proteins in which either the cold-adapted helicase domain (MTM) and/or the N-terminus are exchanged with the respective domains from mesophilic Ded1p. **C.** Change in F350/F330 (left) and back-reflection light scattering (right) as a function of temperature for 4 μM GFP-labelled *S. cerevisiae* Ded1p, *S. kudriavzevii* Ded1p and chimeric proteins (panel B). The apparent T_onset_ are highlighted. Mean (line), sd (light ribbon), n = 6.

**Supplemental Figure 6.**
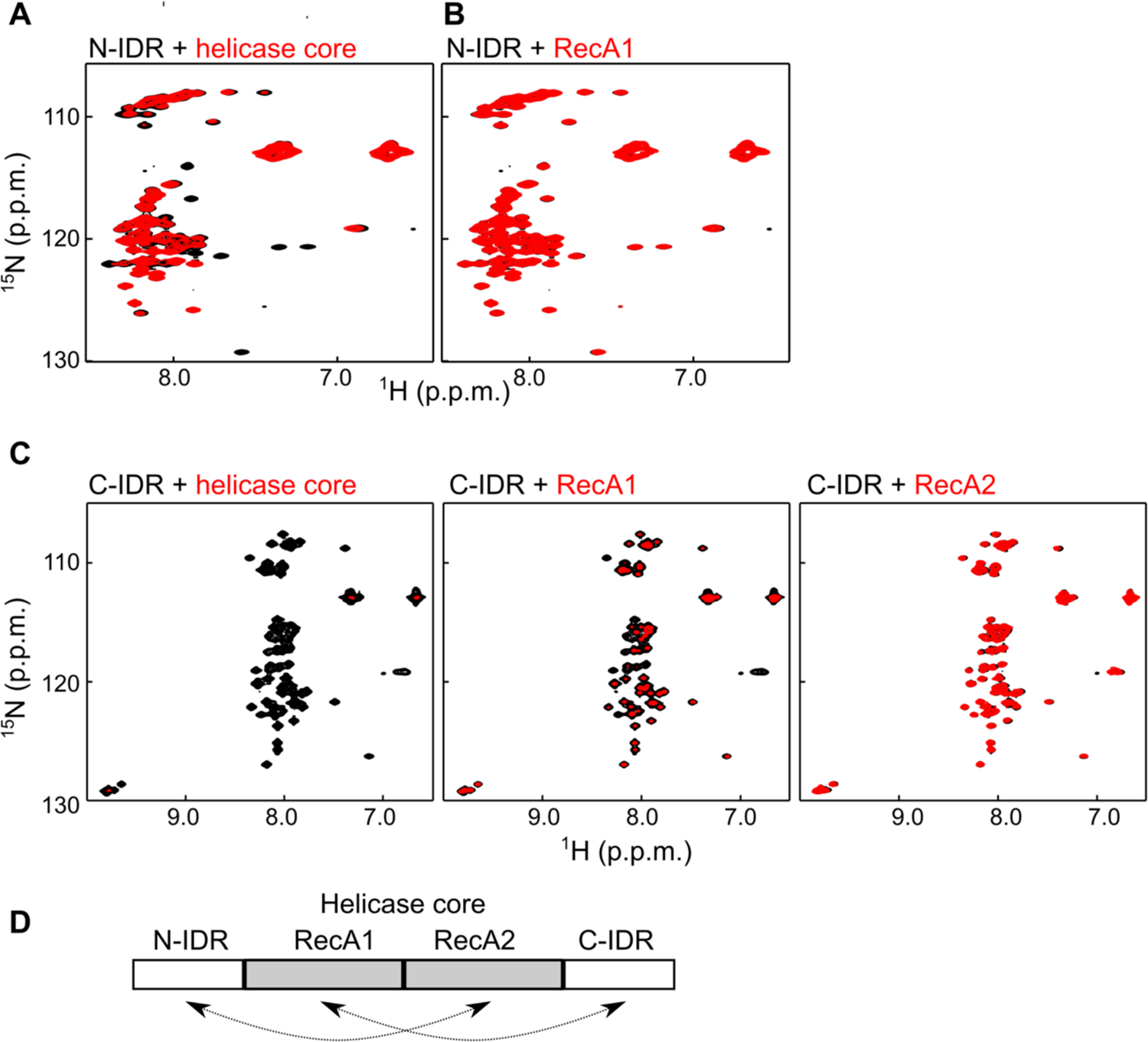
The IDRs interact with the helicase domain. **A.** ^1^H-^15^N NMR spectrum of the N-terminal IDR of Ded1p in the absence (black) and presence (red) of the Ded1p helicase core (RecA1-RecA2) domain. Several the Ded1p resonances experience chemical shift perturbations (CSPs) and broaden upon interaction with the helicase core, indicating a specific binding event. **B**. ^1^H-^15^N NMR spectrum of the N-terminal IDR of Ded1p in the absence (black) and presence (red) of the Ded1p RecA1 domain. The lack of CSPs indicates that the RecA1 domain does not interact with the N-terminal IDR. **C.** ^1^H-^15^N NMR spectra of the C-terminal IDR of Ded1p in the absence (black) and presence (red) of different Ded1p fragments. Left: addition of the helicase core results in severe line broadening, indicating that the C-terminal IDR directly interacts with the core. Middle, addition of the RecA1 domain also results in severe line broadening of the C-IDR resonances, indicating that the C-IDR directly interacts with the C-IDR. Right: the addition of the RecA2 domain to the C-terminal IDR does not result in CSPs, indicating that the C-terminal IDR does not interact with the RecA2 domain. **D.** Summary of the interactions between the Ded1p helical core (RecA1-RecA2) with the N- and C-terminal IDRs.

**Supplemental Figure 7.**
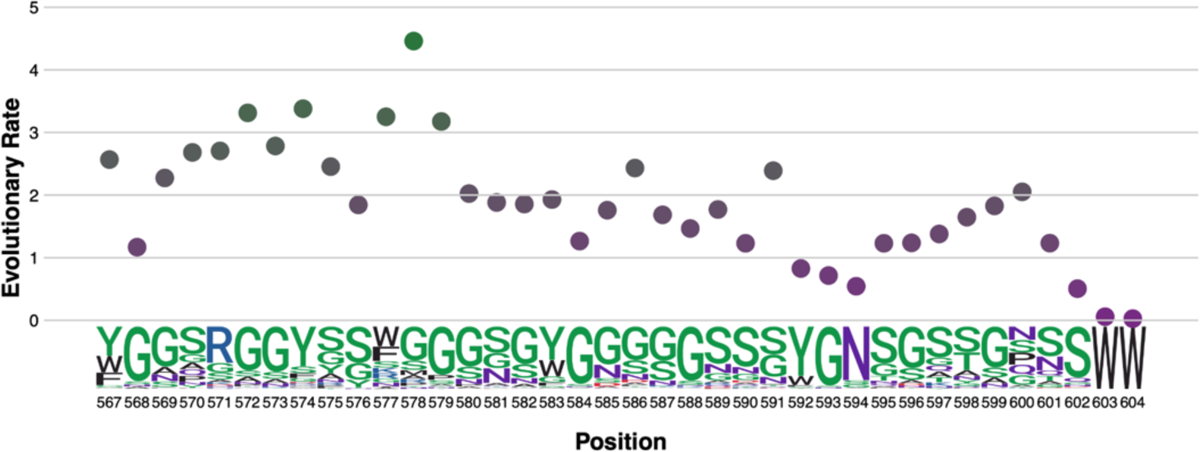
The evolutionary rate calculated for every amino acid in the *Sc* Ded1p C-terminal IDR from position 567-604. Sequence logos and amino acid position are shown below.

**Supplemental Figure 8.**
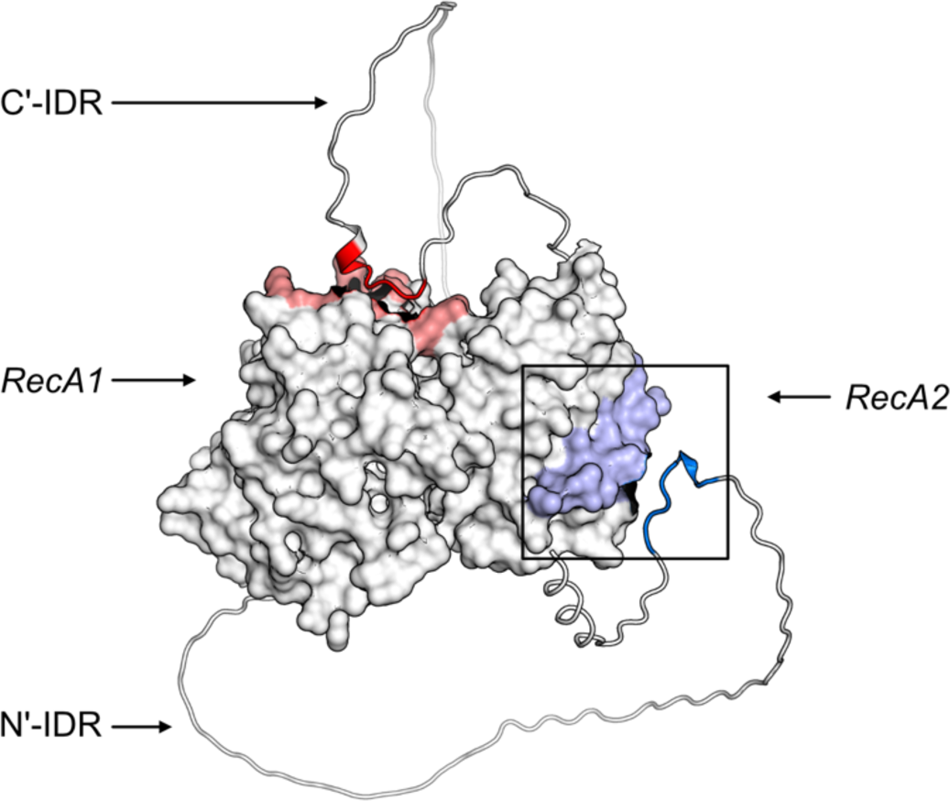
AlphaFold2 model for *Sc* Ded1p (Robinson, 2022; Varadi et al., 2022) highlighting the interaction site between the N-terminal IDR and RecA2 (confirmed with NMR) and the predicted interaction site between the C-terminal IDR and RecA1.

## References

1. Ashwini, C. A review on Chaetomium globosum is versatile weapons for various plant pathogens.

2. Buchan, D.W.A., and Jones, D.T. (2019). The PSIPRED Protein Analysis Workbench: 20 years on. Nucleic Acids Res. 47, W402–W407..

3. Cappelletti, V., Hauser, T., Piazza, I., Pepelnjak, M., Malinovska, L., Fuhrer, T., Li, Y., Dörig, C., Boersema, P., Gillet, L., et al. (2021). Dynamic 3D proteomes reveal protein functional alterations at high resolution in situ. Cell 184, 545–559.e22..

4. Cherkasov, V., Hofmann, S., Druffel-Augustin, S., Mogk, A., Tyedmers, J., Stoecklin, G., and Bukau, B. (2013). Coordination of Translational Control and Protein Homeostasis during Severe Heat Stress. Curr. Biol. 23, 2452–2462..

5. Cherkasov, V., Grousl, T., Theer, P., Vainshtein, Y., Gläßer, C., Mongis, C., Kramer, G., Stoecklin, G., Knop, M., Mogk, A., et al. (2015). Systemic control of protein synthesis through sequestration of translation and ribosome biogenesis factors during severe heat stress. FEBS Lett. 589, 3654–3664..

6. Delaglio, F., Grzesiek, S., Vuister, G.W., Zhu, G., Pfeifer, J., and Bax, A. (1995). NMRPipe: a multidimensional spectral processing system based on UNIX pipes. J. Biomol. NMR 6, 277–293..

7. Dosztányi, Z., Csizmók, V., Tompa, P., and Simon, I. (2005). The pairwise energy content estimated from amino acid composition discriminates between folded and intrinsically unstructured proteins. J. Mol. Biol. 347, 827–839..

8. Epling, L.B., Grace, C.R., Lowe, B.R., Partridge, J.F., and Enemark, E.J. (2015). Cancer-associated mutants of RNA helicase DDX3X are defective in RNA-stimulated ATP hydrolysis. J. Mol. Biol. 427, 1779–1796..

9. Erdős, G., and Dosztányi, Z. (2020). Analyzing protein disorder with IUPred2A. Curr. Protoc. Bioinformatics 70, e99.

10. Fields, P.A. (2001). Review: Protein function at thermal extremes: balancing stability and flexibility. Comp. Biochem. Physiol. A Mol. Integr. Physiol. 129, 417–431..

11. Fields, P.A., Dong, Y., Meng, X., and Somero, G.N. (2015). Adaptations of protein structure and function to temperature: there is more than one way to “skin a cat.” J. Exp. Biol. 218, 1801–1811..

12. Floor, S.N., Condon, K.J., Sharma, D., Jankowsky, E., and Doudna, J.A. (2016). Autoinhibitory interdomain interactions and subfamily-specific extensions redefine the catalytic core of the human DEAD-box protein DDX3. J. Biol. Chem. 291, 2412–2421..

13. Franzmann, T.M., and Alberti, S. (2019). Prion-like low-complexity sequences: Key regulators of protein solubility and phase behavior. J. Biol. Chem. 294, 7128–7136..

14. Franzmann, T.M., Jahnel, M., Pozniakovsky, A., Mahamid, J., Holehouse, A.S., Nüske, E., Richter, D., Baumeister, W., Grill, S.W., Pappu, R.V., et al. (2018). Phase separation of a yeast prion protein promotes cellular fitness. Science 359. https://doi.org/10.1126/science.aao5654.

15. Gao, Z., Putnam, A.A., Bowers, H.A., Guenther, U.-P., Ye, X., Kindsfather, A., Hilliker, A.K., and Jankowsky, E. (2016). Coupling between the DEAD-box RNA helicases Ded1p and eIF4A. Elife 5. https://doi.org/10.7554/eLife.16408.

16. Guerra, E., Chye, P.P., Berardi, E., and Piper, P.W. (2005). Hypoxia abolishes transience of the heat-shock response in the methylotrophic yeast Hansenula polymorpha. Microbiology 151, 805–811..

17. Gulay, S., Gupta, N., Lorsch, J.R., and Hinnebusch, A.G. (2020). Distinct interactions of eIF4A and eIF4E with RNA helicase Ded1 stimulate translation in vivo. Elife 9. https://doi.org/10.7554/eLife.58243.

18. Harmon, T.S., Holehouse, A.S., Rosen, M.K., and Pappu, R.V. (2017). Intrinsically disordered linkers determine the interplay between phase separation and gelation in multivalent proteins. Elife 6. https://doi.org/10.7554/eLife.30294.

19. Hilliker, A., Gao, Z., Jankowsky, E., and Parker, R. (2011). The DEAD-box protein Ded1 modulates translation by the formation and resolution of an eIF4F-mRNA complex. Mol. Cell 43, 962–972..

20. Hoang, D.T., Chernomor, O., von Haeseler, A., Minh, B.Q., and Vinh, L.S. (2018). UFBoot2: Improving the ultrafast bootstrap approximation. Mol. Biol. Evol. 35, 518–522..

21. Högbom, M., Collins, R., van den Berg, S., Jenvert, R.-M., Karlberg, T., Kotenyova, T., Flores, A., Karlsson Hedestam, G.B., and Schiavone, L.H. (2007). Crystal structure of conserved domains 1 and 2 of the human DEAD-box helicase DDX3X in complex with the mononucleotide AMP. J. Mol. Biol. 372, 150–159..

22. Iost, I., Dreyfus, M., and Linder, P. (1999). Ded1p, a DEAD-box protein required for translation initiation in Saccharomyces cerevisiae, is an RNA helicase. J. Biol. Chem. 274, 17677–17683.

23. Iserman, C., Altamirano, C.D., Jegers, C., Friedrich, U., Zarin, T., Fritsch, A.W., Mittasch, M., Domingues, A., Hersemann, L., Jahnel, M., et al. (2020). Condensation of Ded1p Promotes a Translational Switch from Housekeeping to Stress Protein Production. Cell https://doi.org/10.1016/j.cell.2020.04.009.

24. Jarzab, A., Kurzawa, N., Hopf, T., Moerch, M., Zecha, J., Leijten, N., Bian, Y., Musiol, E., Maschberger, M., Stoehr, G., et al. (2020). Meltome atlas-thermal proteome stability across the tree of life. Nat. Methods https://doi.org/10.1038/s41592-020-0801-4.

25. Jumper, J., Evans, R., Pritzel, A., Green, T., Figurnov, M., Ronneberger, O., Tunyasuvunakool, K., Bates, R., Žídek, A., Potapenko, A., et al. (2021). Highly accurate protein structure prediction with AlphaFold. Nature 596, 583–589..

26. Jung, J.-H., Barbosa, A.D., Hutin, S., Kumita, J.R., Gao, M., Derwort, D., Silva, C.S., Lai, X., Pierre, E., Geng, F., et al. (2020). A prion-like domain in ELF3 functions as a thermosensor in Arabidopsis. Nature 585, 256–260..

27. Kroschwald, S., Munder, M.C., Maharana, S., Franzmann, T.M., Richter, D., Ruer, M., Hyman, A.A., and Alberti, S. (2018). Different Material States of Pub1 Condensates Define Distinct Modes of Stress Adaptation and Recovery. Cell Rep. 23, 3327–3339..

28. Lawrence, A.-M., and Besir, H.U.S. (2009). Staining of proteins in gels with Coomassie G-250 without organic solvent and acetic acid. J. Vis. Exp. https://doi.org/10.3791/1350.

29. Lawson Handley, L., Read, D.S., Winfield, I.J., Kimbell, H., Johnson, H., Li, J., Hahn, C., Blackman, R., Wilcox, R., Donnelly, R., et al. (2019). Temporal and spatial variation in distribution of fish environmental DNA in England’s largest lake. Environmental DNA 1, 26–39..

30. Le, S.Q., and Gascuel, O. (2008). An improved general amino acid replacement matrix. Mol. Biol. Evol. 25, 1307–1320..

31. Lemaitre, R.P., Bogdanova, A., Borgonovo, B., Woodruff, J.B., and Drechsel, D.N. (2019). FlexiBAC: a versatile, open-source baculovirus vector system for protein expression, secretion, and proteolytic processing. BMC Biotechnol. 19, 20..

32. Leuenberger, P., Ganscha, S., Kahraman, A., Cappelletti, V., Boersema, P.J., von Mering, C., Claassen, M., and Picotti, P. (2017). Cell-wide analysis of protein thermal unfolding reveals determinants of thermostability. Science 355. https://doi.org/10.1126/science.aai7825.

33. Linder, P., and Jankowsky, E. (2011). From unwinding to clamping - the DEAD box RNA helicase family. Nat. Rev. Mol. Cell Biol. 12, 505–516..

34. Lindquist, S. (1986). The heat-shock response. Annu. Rev. Biochem. 55, 1151–1191..

35. Lindquist, S., and Craig, E.A. (1988). The heat-shock proteins. Annu. Rev. Genet. 22, 631– 677..

36. Mészáros, B., Erdos, G., and Dosztányi, Z. (2018). IUPred2A: context-dependent prediction of protein disorder as a function of redox state and protein binding. Nucleic Acids Res. 46, W329–W337..

37. Minh, B.Q., Schmidt, H.A., Chernomor, O., Schrempf, D., Woodhams, M.D., von Haeseler, A., and Lanfear, R. (2020). IQ-TREE 2: New models and efficient methods for phylogenetic inference in the genomic era. Mol. Biol. Evol. 37, 1530–1534..

38. Morgenstern, I., Powlowski, J., Ishmael, N., Darmond, C., Marqueteau, S., Moisan, M.-C., Quenneville, G., and Tsang, A. (2012). A molecular phylogeny of thermophilic fungi. Fungal Biol. 116, 489–502..

39. van Noort, V., Bradatsch, B., Arumugam, M., Amlacher, S., Bange, G., Creevey, C., Falk, S., Mende, D.R., Sinning, I., Hurt, E., et al. (2013). Consistent mutational paths predict eukaryotic thermostability. BMC Evol. Biol. 13, 7..

40. Pervushin, K., Riek, R., Wider, G., and Wüthrich, K. (1997). Attenuated T2 relaxation by mutual cancellation of dipole-dipole coupling and chemical shift anisotropy indicates an avenue to NMR structures of very large biological macromolecules in solution. Proc. Natl. Acad. Sci. U. S. A. 94, 12366–12371..

41. Riback, J.A., Katanski, C.D., Kear-Scott, J.L., Pilipenko, E.V., Rojek, A.E., Sosnick, T.R., and Drummond, D.A. (2017). Stress-Triggered Phase Separation Is an Adaptive, Evolutionarily Tuned Response. Cell 168, 1028–1040.e19..

42. Robinson, S.L. (2022). Artificial intelligence for microbial biotechnology: beyond the hype. Microb. Biotechnol. 15, 65–69..

43. Ruff, K.M., Choi, Y.H., Cox, D., Ormsby, A.R., Myung, Y., Ascher, D.B., Radford, S.E., Pappu, R.V., and Hatters, D.M. (2022). Sequence grammar underlying the unfolding and phase separation of globular proteins. Mol. Cell https://doi.org/10.1016/j.molcel.2022.06.024.

44. Salvado, Z., Arroyo-Lopez, F.N., Guillamon, J.M., Salazar, G., Querol, A., and Barrio, E. (2011). Temperature Adaptation Markedly Determines Evolution within the Genus Saccharomyces. A PPLIED AND E NVIRONMENTAL M ICROBIOLOGY 77, 2292–2302..

45. Scannell, D.R., Zill, O.A., Rokas, A., Payen, C., Dunham, M.J., Eisen, M.B., Rine, J., Johnston, M., and Hittinger, C.T. (2011). The awesome power of yeast evolutionary genetics: New genome sequences and strain resources for the Saccharomyces sensu stricto genus. G3 (Bethesda) 1, 11–25..

46. Schindelin, J., Arganda-Carreras, I., Frise, E., Kaynig, V., Longair, M., Pietzsch, T., Preibisch, S., Rueden, C., Saalfeld, S., Schmid, B., et al. (2012). Fiji: an open-source platform for biological-image analysis. Nat. Methods 9, 676–682..

47. Sen, N.D., Zhang, H., and Hinnebusch, A.G. (2021). Down-regulation of yeast helicase Ded1 by glucose starvation or heat-shock differentially impairs translation of Ded1-dependent mRNAs. Microorganisms 9, 2413..

48. Sengoku, T., Nureki, O., Nakamura, A., Kobayashi, S., and Yokoyama, S. (2006). Structural basis for RNA unwinding by the DEAD-box protein Drosophila Vasa. Cell 125, 287–300..

49. Sharma, D., and Jankowsky, E. (2014). The Ded1/DDX3 subfamily of DEAD-box RNA helicases. Crit. Rev. Biochem. Mol. Biol. 49, 343–360..

50. Shen, X.-X., Opulente, D.A., Kominek, J., Zhou, X., Steenwyk, J.L., Buh, K.V., Haase, M.A.B., Wisecaver, J.H., Wang, M., Doering, D.T., et al. (2018). Tempo and mode of genome evolution in the budding yeast subphylum. Cell 175, 1533–1545.e20..

51. Somero, G.N. (1995). Proteins and temperature. Annu. Rev. Physiol. 57, 43–68..

52. Song, H., and Ji, X. (2019). The mechanism of RNA duplex recognition and unwinding by DEAD-box helicase DDX3X. Nat. Commun. 10, 3085..

53. Soubrier, J., Steel, M., Lee, M.S.Y., Der Sarkissian, C., Guindon, S., Ho, S.Y.W., and Cooper, A. (2012). The influence of rate heterogeneity among sites on the time dependence of molecular rates. Mol. Biol. Evol. 29, 3345–3358..

54. Tabari, E., and Su, Z. (2017). PorthoMCL: Parallel orthology prediction using MCL for the realm of massive genome availability. Big Data Anal. 2. https://doi.org/10.1186/s41044-016-0019-8.

55. Taylor, T.J., and Vaisman, I.I. (2010). Discrimination of thermophilic and mesophilic proteins. BMC Struct. Biol. 10 *Suppl 1*, S5..

56. UniProt Consortium (2019). UniProt: a worldwide hub of protein knowledge. Nucleic Acids Res. 47, D506–D515..

57. Varadi, M., Anyango, S., Deshpande, M., Nair, S., Natassia, C., Yordanova, G., Yuan, D., Stroe, O., Wood, G., Laydon, A., et al. (2022). AlphaFold Protein Structure Database: massively expanding the structural coverage of protein-sequence space with high-accuracy models. Nucleic Acids Res. 50, D439–D444..

58. Verghese, J., Abrams, J., Wang, Y., and Morano, K.A. (2012). Biology of the heat shock response and protein chaperones: budding yeast (Saccharomyces cerevisiae) as a model system. Microbiol. Mol. Biol. Rev. 76, 115–158..

59. Vijayakumar, J., Perrois, C., Heim, M., Bousset, L., Alberti, S., and Besse, F. (2019). The prion-like domain of Drosophila Imp promotes axonal transport of RNP granules in vivo. Nat. Commun. 10, 2593..

60. Wallace, E.W.J., Kear-Scott, J.L., Pilipenko, E.V., Schwartz, M.H., Laskowski, P.R., Rojek, A.E., Katanski, C.D., Riback, J.A., Dion, M.F., Franks, A.M., et al. (2015). Reversible, Specific, Active Aggregates of Endogenous Proteins Assemble upon Heat Stress. Cell 162, 1286–1298..

61. Waterhouse, A.M., Procter, J.B., Martin, D.M.A., Clamp, M., and Barton, G.J. (2009). Jalview Version 2--a multiple sequence alignment editor and analysis workbench. Bioinformatics 25, 1189–1191..

62. Wilkinson, E.G., and Strader, L.C. (2020). A prion-based thermosensor in plants. Mol. Cell 80, 181–182..

63. Yang, Z. (1995). A space-time process model for the evolution of DNA sequences. Genetics 139, 993–1005..

64. Yoo, H., Bard, J.A.M., Pilipenko, E.V., and Drummond, D.A. (2022). Chaperones directly and efficiently disperse stress-triggered biomolecular condensates. Mol. Cell 82, 741–755.e11..

